## Supplementary material for "Fast label-free live imaging reveals key roles of flow dynamics and CD44-HA interaction in cancer cell arrest on endothelial monolayers": Main figures Statistical summaries

### Figure 1 to figure 8, Statistical summaries

#### Fig1C

| Condition | count | mean | std | min | 25% | 50% | 75% | max |
|---|---|---|---|---|---|---|---|---|
| AsPC1 | 28.0 | 56.286 | 32.863 | 4.0 | 24.0 | 63.0 | 76.5 | 125.0 |
| BxPC3 | 30.0 | 18.1 | 29.169 | 0.0 | 1.0 | 4.5 | 16.75 | 102.0 |
| MiaPaca2 | 36.0 | 63.306 | 45.875 | 7.0 | 28.5 | 47.0 | 99.5 | 180.0 |
| PANC1 | 30.0 | 10.133 | 11.617 | 0.0 | 2.0 | 4.5 | 16.75 | 39.0 |
| Panc10.05 | 32.0 | 16.219 | 14.827 | 0.0 | 2.75 | 17.0 | 23.0 | 62.0 |
| SU86.86 | 25.0 | 16.32 | 18.925 | 1.0 | 6.0 | 11.0 | 21.0 | 95.0 |
| SW1990 | 25.0 | 23.44 | 23.988 | 1.0 | 5.0 | 12.0 | 29.0 | 74.0 |

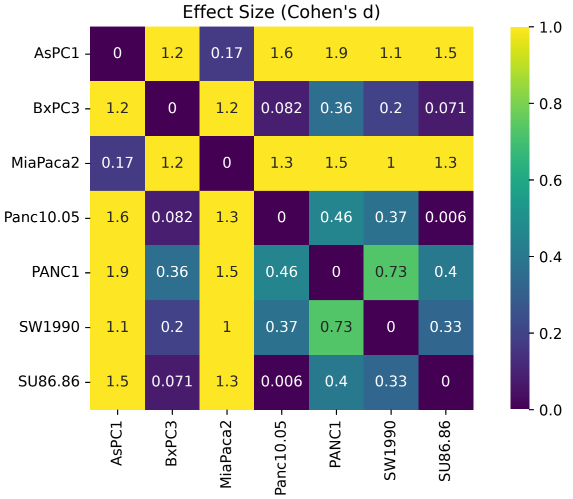


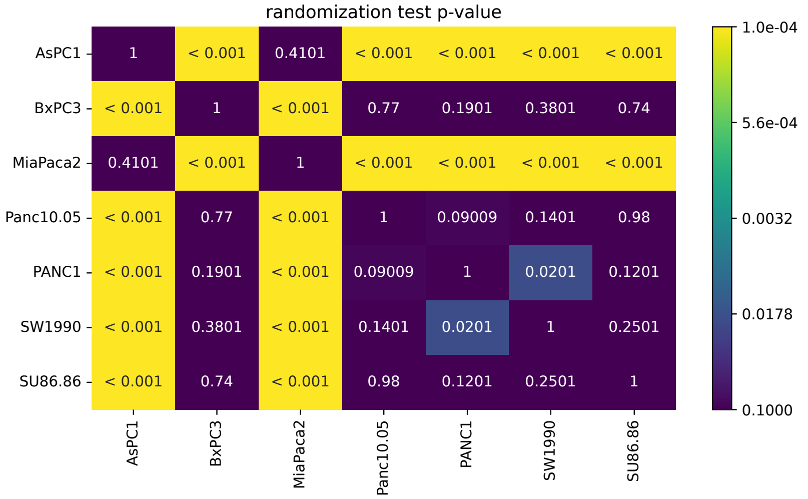


#### Fig1H

| Condition | count | mean | std | min | 25% | 50% | 75% | max |
|---|---|---|---|---|---|---|---|---|
| AsPc1_100_CTRL | 7.0 | 0.232 | 0.264 | 0.067 | 0.088 | 0.111 | 0.23 | 0.809 |
| AsPc1_200_CTRL | 7.0 | 0.09 | 0.054 | 0.031 | 0.057 | 0.073 | 0.109 | 0.195 |
| AsPc1_300_CTRL | 7.0 | 0.035 | 0.046 | -0.013 | 0.0 | 0.038 | 0.048 | 0.121 |
| MiaPaca-2_100_CTRL | 6.0 | 0.349 | 0.45 | -0.198 | 0.093 | 0.237 | 0.635 | 1.008 |
| MiaPaca-2_200_CTRL | 6.0 | 0.419 | 0.305 | -0.042 | 0.279 | 0.409 | 0.633 | 0.794 |
| MiaPaca-2_300_CTRL | 6.0 | 0.385 | 0.415 | 0.034 | 0.17 | 0.242 | 0.407 | 1.183 |
| Panc10_100_CTRL | 4.0 | 0.011 | 0.015 | -0.005 | 0.0 | 0.01 | 0.021 | 0.027 |
| Panc10_200_CTRL | 4.0 | -0.001 | 0.003 | -0.005 | -0.002 | -0.0 | 0.001 | 0.003 |
| Panc10_300_CTRL | 4.0 | -0.0 | 0.021 | -0.03 | -0.005 | 0.005 | 0.008 | 0.019 |

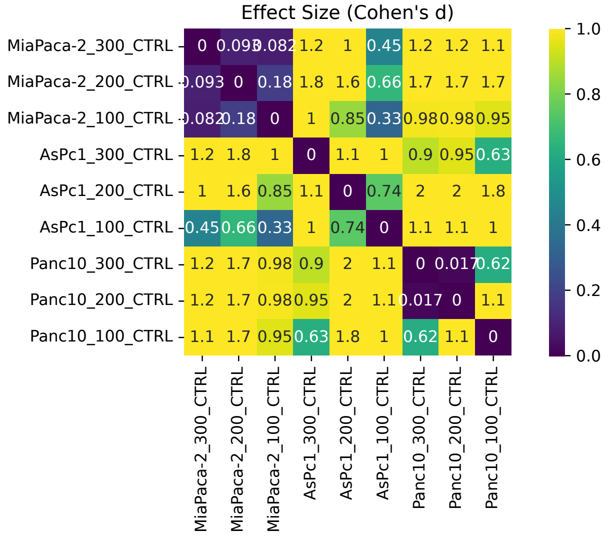


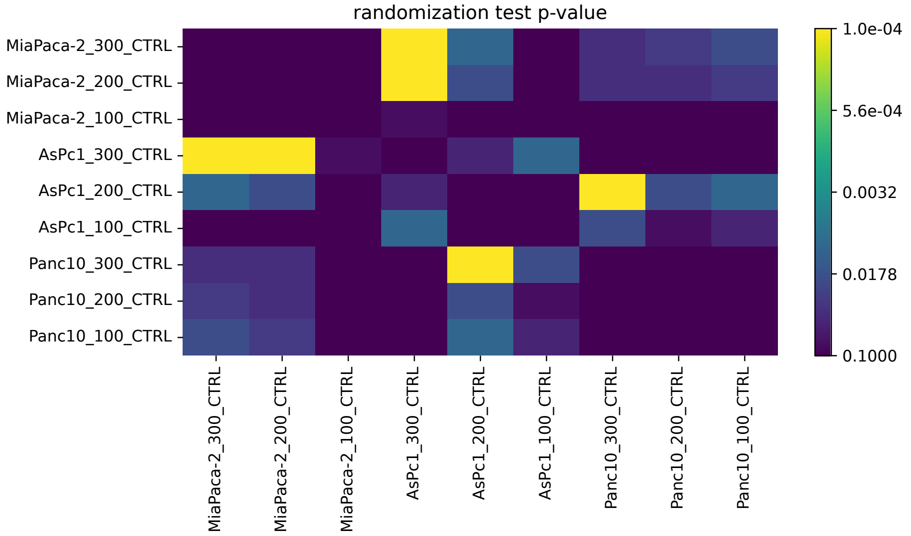

### Fig1J

| Condition | count | mean | std | min | 25% | 50% | 75% | max |
|---|---|---|---|---|---|---|---|---|
| As_100 | 7329.0 | 182.931 | 174.931 | 3.183 | 30.563 | 142.237 | 261.224 | 842.187 |
| As_200 | 7329.0 | 378.023 | 203.595 | 4.193 | 239.853 | 374.344 | 560.912 | 874.253 |
| As_300 | 6854.0 | 429.691 | 185.496 | 3.275 | 296.862 | 443.094 | 614.088 | 726.71 |
| As_wash | 5325.0 | 139.246 | 185.128 | 3.145 | 18.412 | 42.998 | 176.637 | 795.132 |
| Mia_100 | 3010.0 | 87.874 | 99.071 | 2.284 | 14.796 | 42.482 | 150.363 | 689.734 |
| Mia_200 | 3232.0 | 104.429 | 121.029 | 1.624 | 24.33 | 57.613 | 139.641 | 834.089 |
| Mia_300 | 3232.0 | 116.472 | 137.663 | 3.312 | 27.823 | 64.195 | 143.041 | 838.898 |
| Mia_wash | 4732.0 | 64.127 | 76.522 | 2.613 | 15.298 | 32.14 | 86.659 | 868.248 |
| P10_100 | 8226.0 | 383.195 | 205.878 | 5.991 | 195.625 | 344.828 | 631.405 | 828.402 |
| P10_200 | 8226.0 | 462.033 | 178.126 | 7.995 | 320.346 | 477.507 | 652.482 | 747.1 |
| P10_300 | 6857.0 | 493.389 | 151.43 | 4.775 | 371.646 | 511.88 | 650.41 | 747.1 |
| P10_wash | 5935.0 | 541.9 | 130.297 | 8.856 | 449.489 | 596.199 | 653.094 | 758.636 |

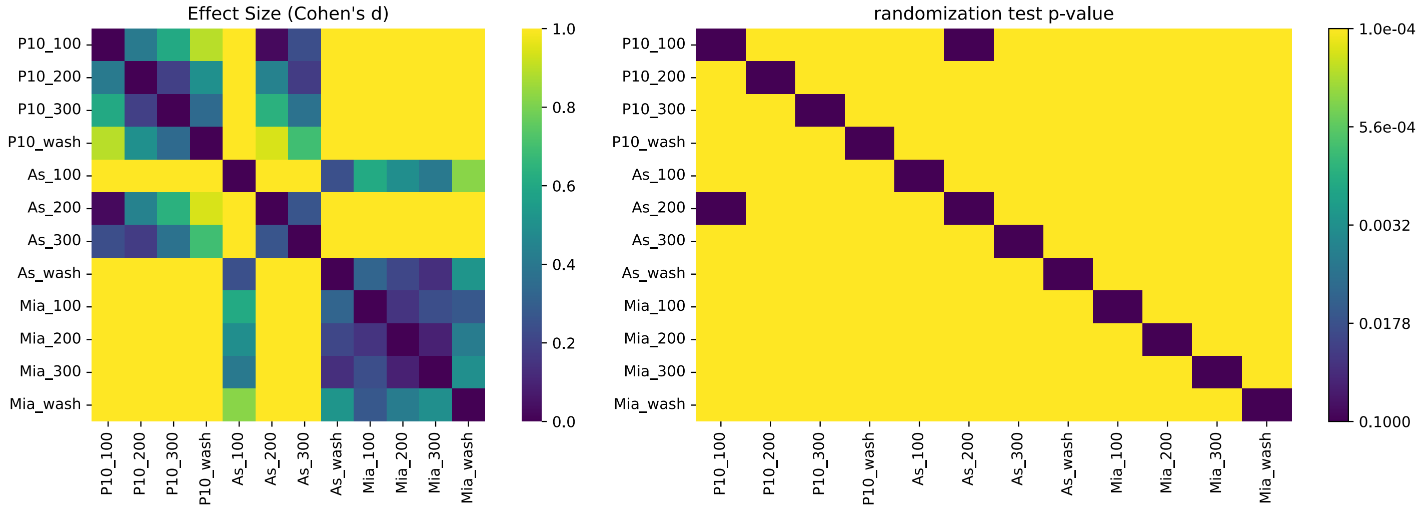


### Fig1K

| Condition | count | mean | std | min | 25% | 50% | 75% | max |
|---|---|---|---|---|---|---|---|---|
| As_100 | 7329.0 | 0.626 | 0.447 | -0.567 | 0.049 | 0.955 | 0.986 | 0.999 |
| As_200 | 7329.0 | 0.858 | 0.322 | -0.361 | 0.973 | 0.994 | 0.997 | 1.0 |
| As_300 | 6854.0 | 0.918 | 0.246 | -0.551 | 0.986 | 0.996 | 0.998 | 1.0 |
| As_wash | 5325.0 | 0.244 | 0.407 | -0.708 | 0.003 | 0.025 | 0.399 | 1.0 |
| Mia_100 | 3010.0 | 0.101 | 0.263 | -0.417 | -0.009 | 0.004 | 0.041 | 0.999 |
| Mia_200 | 3232.0 | 0.195 | 0.307 | -0.485 | 0.002 | 0.041 | 0.267 | 0.999 |
| Mia_300 | 3232.0 | 0.312 | 0.346 | -0.573 | 0.022 | 0.175 | 0.589 | 1.0 |
| Mia_wash | 4732.0 | 0.072 | 0.193 | -0.623 | -0.002 | 0.015 | 0.047 | 0.997 |
| P10_100 | 8226.0 | 0.977 | 0.095 | -0.292 | 0.986 | 0.991 | 0.994 | 0.999 |
| P10_200 | 8226.0 | 0.988 | 0.059 | -0.005 | 0.992 | 0.996 | 0.997 | 1.0 |
| P10_300 | 6857.0 | 0.986 | 0.088 | -0.053 | 0.995 | 0.997 | 0.998 | 1.0 |
| P10_wash | 5935.0 | 0.982 | 0.086 | -0.177 | 0.993 | 0.997 | 0.998 | 1.0 |

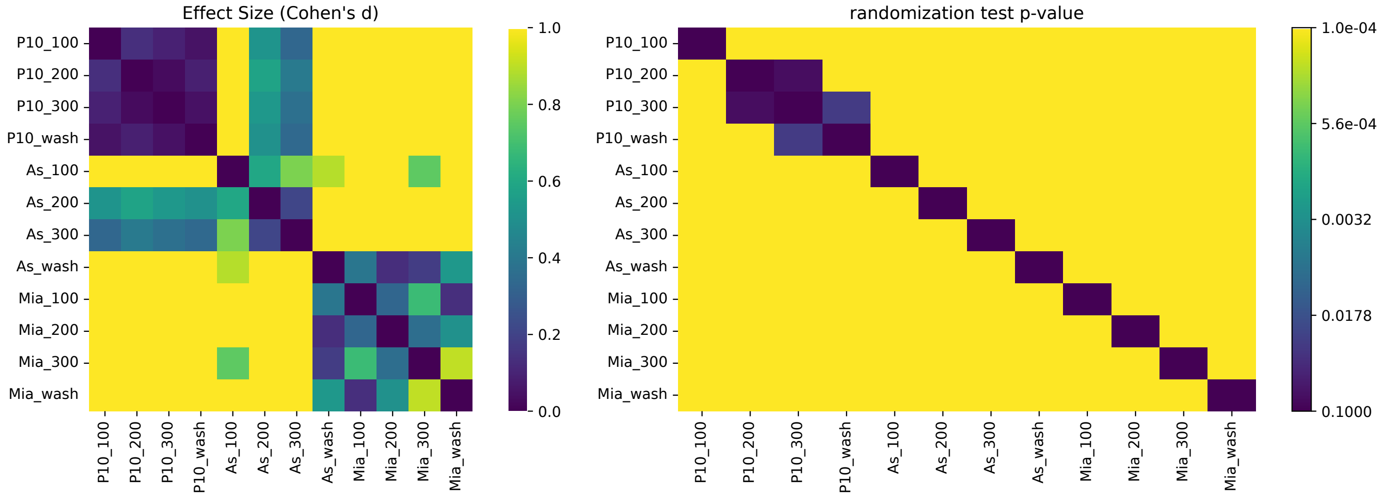

### Fig2D

| Condition | count | mean | std | min | 25% | 50% | 75% | max |
|---|---|---|---|---|---|---|---|---|
| Monocyte_100_CTRL | 6.0 | 0.014 | 0.039 | -0.017 | -0.007 | -0.0 | 0.015 | 0.09 |
| Monocyte_100_IL1b | 4.0 | 0.084 | 0.208 | -0.106 | -0.022 | 0.032 | 0.138 | 0.379 |
| Monocyte_200_CTRL | 6.0 | 0.081 | 0.138 | -0.007 | 0.009 | 0.032 | 0.063 | 0.357 |
| Monocyte_200_IL1b | 4.0 | 0.319 | 0.295 | 0.059 | 0.125 | 0.247 | 0.441 | 0.723 |
| Monocyte_300_CTRL | 6.0 | 0.069 | 0.076 | -0.065 | 0.05 | 0.083 | 0.123 | 0.141 |
| Monocyte_300_IL1b | 4.0 | 0.168 | 0.207 | -0.008 | 0.026 | 0.114 | 0.256 | 0.451 |
| Neutrophil_100_CTRL | 8.0 | 0.572 | 0.308 | 0.164 | 0.327 | 0.53 | 0.863 | 0.965 |
| Neutrophil_100_IL1b | 4.0 | 0.512 | 0.306 | 0.209 | 0.272 | 0.507 | 0.747 | 0.825 |
| Neutrophil_200_CTRL | 8.0 | 0.566 | 0.247 | 0.204 | 0.383 | 0.577 | 0.717 | 0.912 |
| Neutrophil_200_IL1b | 4.0 | 0.76 | 0.627 | 0.19 | 0.297 | 0.648 | 1.111 | 1.554 |
| Neutrophil_300_CTRL | 8.0 | 0.35 | 0.205 | 0.057 | 0.201 | 0.424 | 0.52 | 0.537 |
| Neutrophil_300_IL1b | 4.0 | 0.759 | 0.634 | 0.287 | 0.408 | 0.531 | 0.883 | 1.689 |

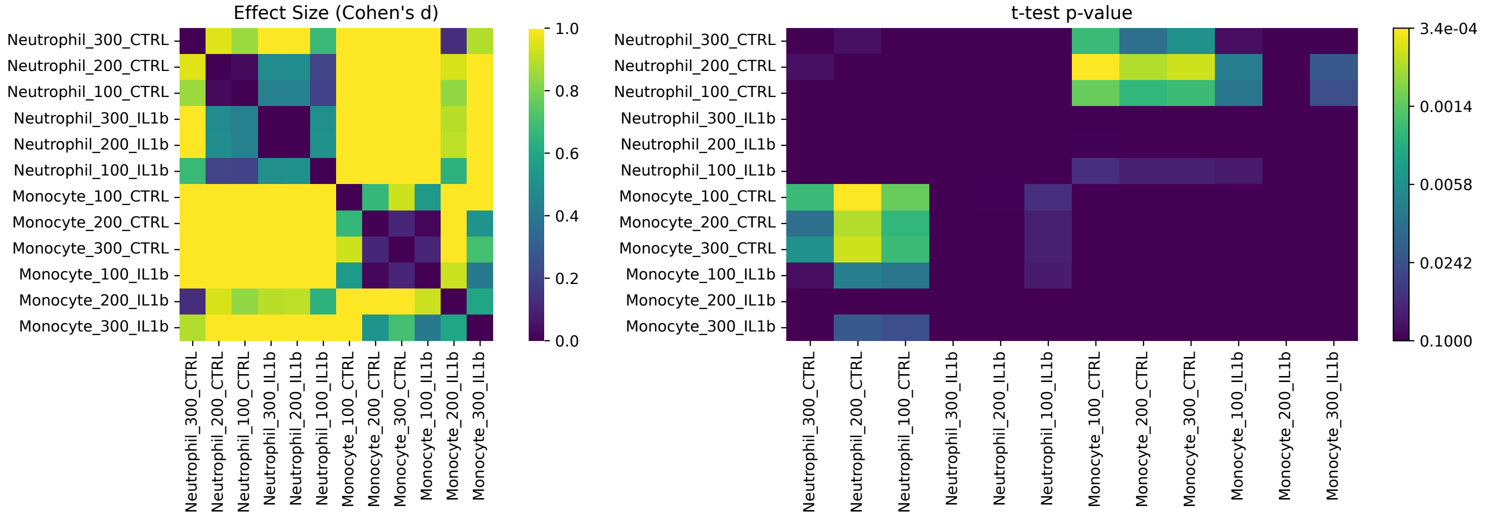


### Fig2F

| Condition | count | mean | std | min | 25% | 50% | 75% | max |
|---|---|---|---|---|---|---|---|---|
| Monocyte_100_CTRL | 6.0 | 0.014 | 0.039 | -0.017 | -0.007 | -0.0 | 0.015 | 0.09 |
| Monocyte_100_IL1b | 4.0 | 0.084 | 0.208 | -0.106 | -0.022 | 0.032 | 0.138 | 0.379 |
| Monocyte_200_CTRL | 6.0 | 0.081 | 0.138 | -0.007 | 0.009 | 0.032 | 0.063 | 0.357 |
| Monocyte_200_IL1b | 4.0 | 0.319 | 0.295 | 0.059 | 0.125 | 0.247 | 0.441 | 0.723 |
| Monocyte_300_CTRL | 6.0 | 0.069 | 0.076 | -0.065 | 0.05 | 0.083 | 0.123 | 0.141 |
| Monocyte_300_IL1b | 4.0 | 0.168 | 0.207 | -0.008 | 0.026 | 0.114 | 0.256 | 0.451 |
| Neutrophil_100_CTRL | 8.0 | 0.572 | 0.308 | 0.164 | 0.327 | 0.53 | 0.863 | 0.965 |
| Neutrophil_100_IL1b | 4.0 | 0.512 | 0.306 | 0.209 | 0.272 | 0.507 | 0.747 | 0.825 |
| Neutrophil_200_CTRL | 8.0 | 0.566 | 0.247 | 0.204 | 0.383 | 0.577 | 0.717 | 0.912 |
| Neutrophil_200_IL1b | 4.0 | 0.76 | 0.627 | 0.19 | 0.297 | 0.648 | 1.111 | 1.554 |
| Neutrophil_300_CTRL | 8.0 | 0.35 | 0.205 | 0.057 | 0.201 | 0.424 | 0.52 | 0.537 |
| Neutrophil_300_IL1b | 4.0 | 0.759 | 0.634 | 0.287 | 0.408 | 0.531 | 0.883 | 1.689 |

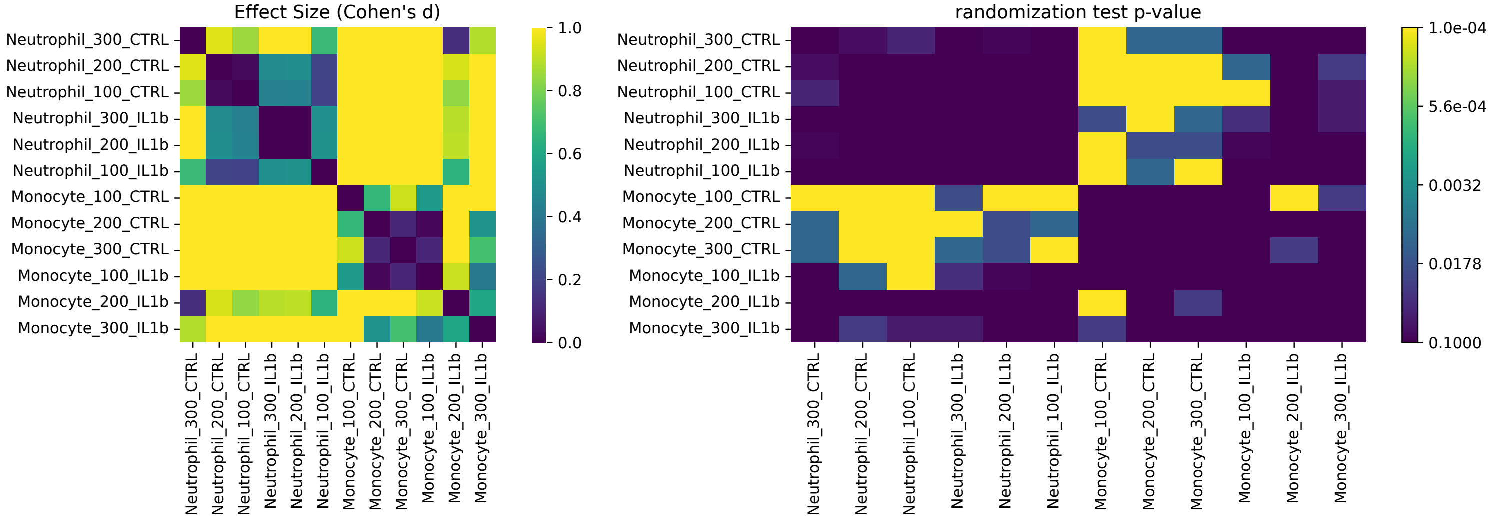

Fig2G

| Condition | count | mean | std | min | 25% | 50% | 75% | max |
|---|---|---|---|---|---|---|---|---|
| IL_Mono_100 | 7031.0 | 0.703 | 0.425 | -0.595 | 0.123 | 0.975 | 0.986 | 0.999 |
| IL_Mono_200 | 7031.0 | 0.744 | 0.404 | -0.556 | 0.621 | 0.986 | 0.995 | 1.0 |
| IL_Mono_300 | 5564.0 | 0.747 | 0.404 | -0.556 | 0.655 | 0.992 | 0.997 | 1.0 |
| IL_Mono_wash | 8188.0 | 0.714 | 0.383 | -0.627 | 0.542 | 0.927 | 0.995 | 1.0 |
| IL_Neut_100 | 4451.0 | 0.259 | 0.393 | -0.567 | -0.003 | 0.03 | 0.623 | 0.998 |
| IL_Neut_200 | 4450.0 | 0.281 | 0.407 | -0.567 | -0.003 | 0.034 | 0.69 | 0.999 |
| IL_Neut_300 | 3044.0 | 0.362 | 0.427 | -0.473 | 0.002 | 0.085 | 0.872 | 1.0 |
| IL_Neut_wash | 4976.0 | 0.171 | 0.332 | -0.622 | -0.004 | 0.021 | 0.15 | 0.999 |
| Mono_100 | 10947.0 | 0.914 | 0.238 | -0.356 | 0.978 | 0.988 | 0.992 | 0.999 |
| Mono_200 | 10947.0 | 0.931 | 0.222 | -0.458 | 0.984 | 0.995 | 0.997 | 1.0 |
| Mono_300 | 8045.0 | 0.906 | 0.271 | -0.657 | 0.989 | 0.996 | 0.998 | 1.0 |
| Mono_wash | 12548.0 | 0.936 | 0.198 | -0.507 | 0.969 | 0.996 | 0.998 | 1.0 |
| Neut_100 | 10063.0 | 0.438 | 0.449 | -0.61 | 0.007 | 0.223 | 0.925 | 0.999 |
| Neut_200 | 10063.0 | 0.496 | 0.464 | -0.591 | 0.01 | 0.551 | 0.987 | 1.0 |
| Neut_300 | 7028.0 | 0.585 | 0.457 | -0.561 | 0.027 | 0.919 | 0.994 | 1.0 |
| Neut_wash | 12904.0 | 0.467 | 0.454 | -0.703 | 0.014 | 0.384 | 0.98 | 1.0 |

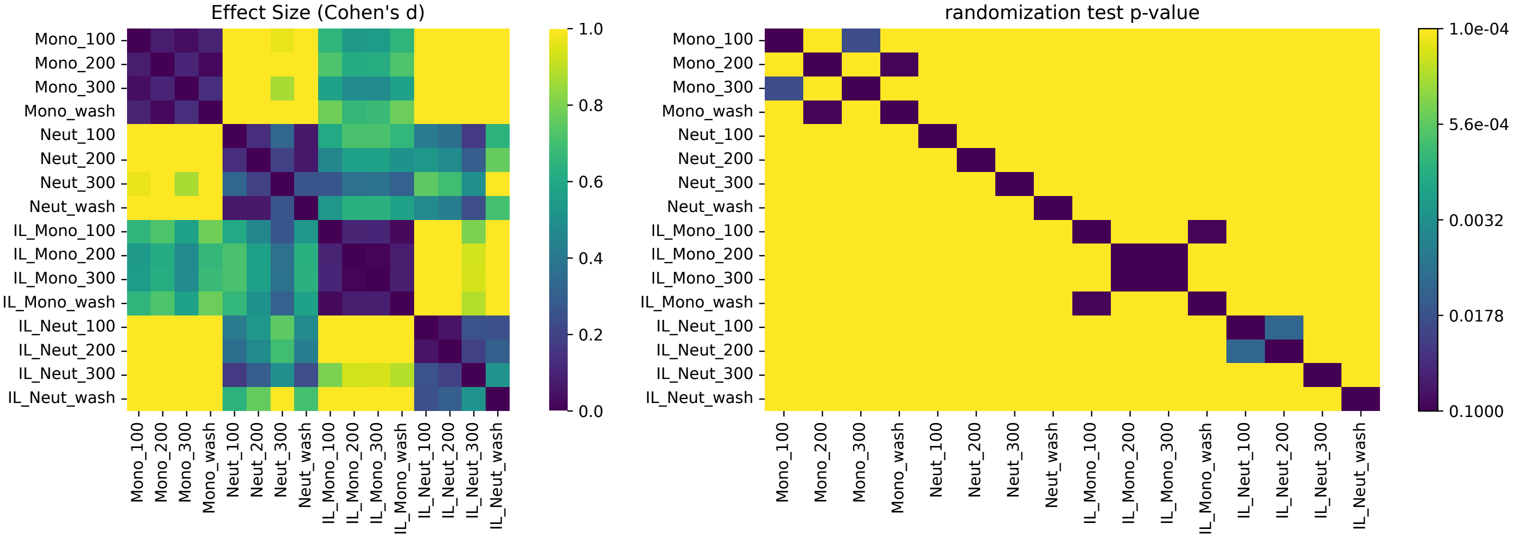

Fig2L

| Condition | count | mean | std | min | 25% | 50% | 75% | max |
|---|---|---|---|---|---|---|---|---|
| AsPc1_100_CTRL | 7.0 | 0.232 | 0.264 | 0.067 | 0.088 | 0.111 | 0.23 | 0.809 |
| AsPc1_100_IL1b | 3.0 | 0.206 | 0.115 | 0.074 | 0.168 | 0.261 | 0.272 | 0.283 |
| AsPc1_200_CTRL | 7.0 | 0.09 | 0.054 | 0.031 | 0.057 | 0.073 | 0.109 | 0.195 |
| AsPc1_200_IL1b | 3.0 | 0.015 | 0.016 | -0.002 | 0.008 | 0.018 | 0.024 | 0.029 |
| AsPc1_300_CTRL | 7.0 | 0.035 | 0.046 | -0.013 | 0.0 | 0.038 | 0.048 | 0.121 |
| AsPc1_300_IL1b | 3.0 | -0.001 | 0.014 | -0.016 | -0.008 | -0.0 | 0.006 | 0.013 |
| MiaPaca-2_100_CTRL | 6.0 | 0.349 | 0.45 | -0.198 | 0.093 | 0.237 | 0.635 | 1.008 |
| MiaPaca-2_100_IL1b | 2.0 | 0.981 | 0.371 | 0.718 | 0.85 | 0.981 | 1.112 | 1.243 |
| MiaPaca-2_200_CTRL | 6.0 | 0.419 | 0.305 | -0.042 | 0.279 | 0.409 | 0.633 | 0.794 |
| MiaPaca-2_200_IL1b | 2.0 | 0.735 | 0.26 | 0.551 | 0.643 | 0.735 | 0.827 | 0.918 |
| MiaPaca-2_300_CTRL | 6.0 | 0.385 | 0.415 | 0.034 | 0.17 | 0.242 | 0.407 | 1.183 |
| MiaPaca-2_300_IL1b | 2.0 | 0.494 | 0.046 | 0.461 | 0.477 | 0.494 | 0.51 | 0.526 |
| Monocyte_100_CTRL | 6.0 | 0.014 | 0.039 | -0.017 | -0.007 | -0.0 | 0.015 | 0.09 |
| Monocyte_100_IL1b | 4.0 | 0.084 | 0.208 | -0.106 | -0.022 | 0.032 | 0.138 | 0.379 |
| Monocyte_200_CTRL | 6.0 | 0.081 | 0.138 | -0.007 | 0.009 | 0.032 | 0.063 | 0.357 |
| Monocyte_200_IL1b | 4.0 | 0.319 | 0.295 | 0.059 | 0.125 | 0.247 | 0.441 | 0.723 |
| Monocyte_300_CTRL | 6.0 | 0.069 | 0.076 | -0.065 | 0.05 | 0.083 | 0.123 | 0.141 |
| Monocyte_300_IL1b | 4.0 | 0.168 | 0.207 | -0.008 | 0.026 | 0.114 | 0.256 | 0.451 |
| Neutrophil_100_CTRL | 8.0 | 0.572 | 0.308 | 0.164 | 0.327 | 0.53 | 0.863 | 0.965 |
| Neutrophil_100_IL1b | 4.0 | 0.512 | 0.306 | 0.209 | 0.272 | 0.507 | 0.747 | 0.825 |
| Neutrophil_200_CTRL | 8.0 | 0.566 | 0.247 | 0.204 | 0.383 | 0.577 | 0.717 | 0.912 |
| Neutrophil_200_IL1b | 4.0 | 0.76 | 0.627 | 0.19 | 0.297 | 0.648 | 1.111 | 1.554 |
| Neutrophil_300_CTRL | 8.0 | 0.35 | 0.205 | 0.057 | 0.201 | 0.424 | 0.52 | 0.537 |
| Neutrophil_300_IL1b | 4.0 | 0.759 | 0.634 | 0.287 | 0.408 | 0.531 | 0.883 | 1.689 |
| Panc10_100_CTRL | 4.0 | 0.011 | 0.015 | -0.005 | 0.0 | 0.01 | 0.021 | 0.027 |
| Panc10_100_IL1b | 4.0 | 0.12 | 0.143 | 0.001 | 0.053 | 0.076 | 0.144 | 0.327 |
| Panc10_200_CTRL | 4.0 | -0.001 | 0.003 | -0.005 | -0.002 | -0.0 | 0.001 | 0.003 |
| Panc10_200_IL1b | 4.0 | 0.026 | 0.045 | -0.033 | 0.007 | 0.031 | 0.049 | 0.074 |
| Panc10_300_CTRL | 4.0 | -0.0 | 0.021 | -0.03 | -0.005 | 0.005 | 0.008 | 0.019 |
| Panc10_300_IL1b | 4.0 | 0.006 | 0.014 | -0.012 | -0.002 | 0.009 | 0.017 | 0.019 |

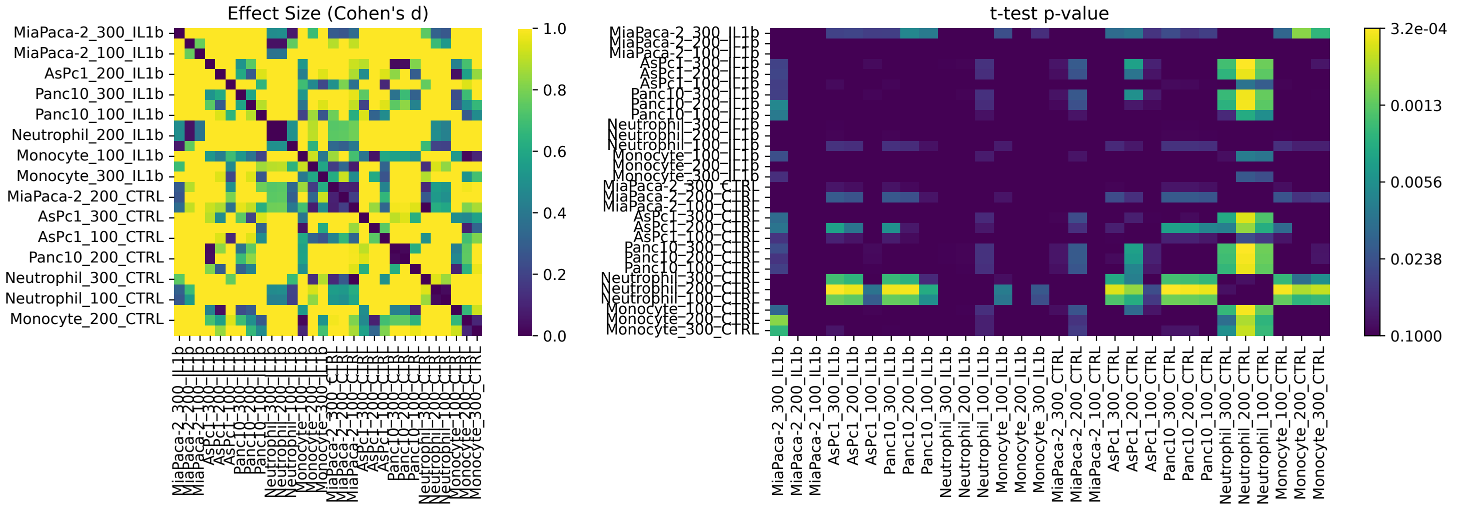

Fig3C

| Condition | count | mean | std | min | 25% | 50% | 75% | max |
|---|---|---|---|---|---|---|---|---|
| AsPC1_Ctrl | 334.0 | 41.822 | 86.576 | 0.0 | 1.626 | 7.908 | 37.961 | 573.84 |
| AsPC1_IL1b | 327.0 | 44.67 | 89.951 | 0.0 | 1.648 | 7.687 | 39.616 | 573.84 |
| MiaPaca2_IL1b | 393.0 | 36.641 | 59.326 | 0.0 | 5.621 | 17.102 | 39.011 | 471.969 |
| MiaPaca2_ Ctrl | 395.0 | 31.4 | 44.534 | 0.0 | 6.237 | 17.406 | 37.417 | 413.337 |
| Mono._Ctrl | 1116.0 | 20.379 | 53.656 | 0.0 | 1.322 | 4.476 | 14.053 | 557.169 |
| Mono._IL1b | 1105.0 | 21.399 | 58.974 | 0.0 | 0.798 | 2.842 | 11.582 | 558.456 |
| Neutro._Ctrl | 1710.0 | 38.964 | 82.03 | 0.0 | 2.539 | 8.542 | 26.988 | 643.54 |
| Neutro._IL1b | 1692.0 | 29.037 | 59.493 | 0.0 | 2.154 | 6.617 | 24.192 | 583.495 |

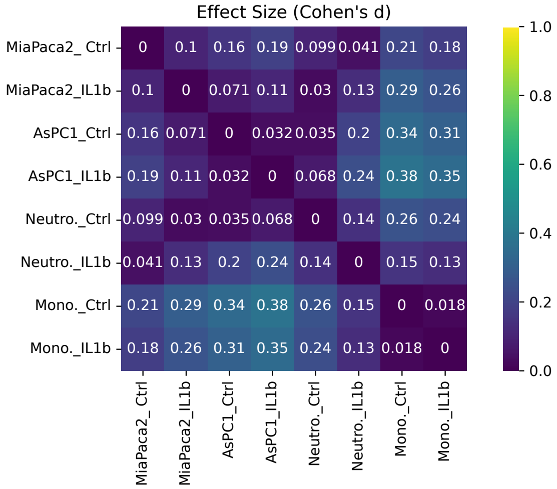


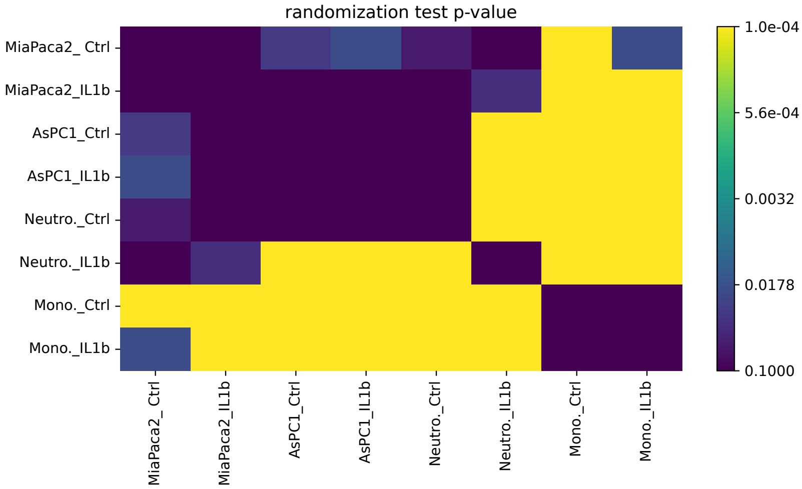


Fig3D

| Condition | count | mean | std | min | 25% | 50% | 75% | max |
|---|---|---|---|---|---|---|---|---|
| AsPC1_Ctrl | 335.0 | 49.597 | 70.279 | 0.0 | 5.333 | 17.228 | 66.806 | 460.171 |
| AsPC1_IL | 335.0 | 49.544 | 71.723 | 0.092 | 5.451 | 19.302 | 67.338 | 460.171 |
| MiaPaca2_Ctrl | 395.0 | 110.223 | 118.49 | 0.0 | 19.889 | 59.087 | 165.148 | 596.886 |
| MiaPaca2_IL | 394.0 | 98.938 | 111.059 | 0.0 | 18.072 | 53.356 | 147.42 | 596.886 |
| Mono_Ctrl | 1138.0 | 67.798 | 84.834 | 0.0 | 8.985 | 35.103 | 92.311 | 695.421 |
| Mono_IL | 1105.0 | 36.207 | 54.971 | 0.0 | 5.339 | 16.3 | 43.435 | 454.098 |
| Neutro_IL | 1708.0 | 87.441 | 92.361 | 0.0 | 19.853 | 57.669 | 128.523 | 724.838 |
| Neutro_ctrl | 1710.0 | 90.143 | 93.212 | 0.0 | 18.624 | 60.397 | 136.562 | 749.723 |

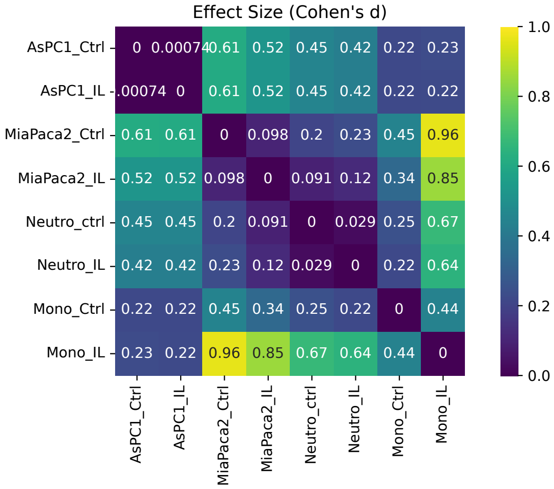


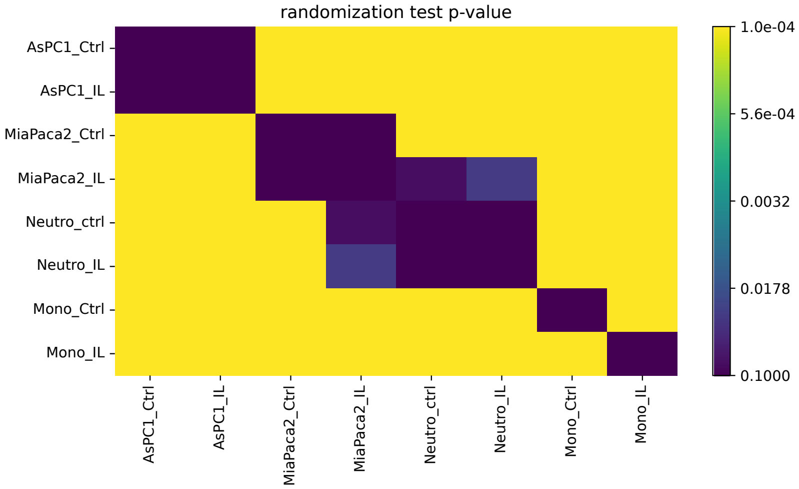

#### Fig3E

| Condition | count | mean | std | min | 25% | 50% | 75% | max |
|---|---|---|---|---|---|---|---|---|
| AsPC1_Ctrl | 335.0 | 0.105 | 0.241 | -0.705 | -0.0 | 0.024 | 0.102 | 0.965 |
| AsPC1_IL | 335.0 | 0.086 | 0.228 | -0.705 | -0.002 | 0.02 | 0.091 | 0.965 |
| MiaPaca2_Ctrl | 394.0 | 0.128 | 0.205 | -0.854 | 0.012 | 0.067 | 0.187 | 0.97 |
| MiaPaca2_IL | 393.0 | 0.089 | 0.194 | -0.998 | 0.007 | 0.047 | 0.142 | 0.855 |
| Mono_Ctrl | 1167.0 | 0.046 | 0.168 | -0.676 | -0.01 | 0.009 | 0.047 | 0.998 |
| Mono_IL | 1106.0 | 0.092 | 0.222 | -0.934 | -0.004 | 0.015 | 0.093 | 0.993 |
| Neutro_IL | 1708.0 | 0.059 | 0.18 | -0.669 | -0.009 | 0.01 | 0.055 | 0.998 |
| Neutro_ctrl | 1710.0 | 0.053 | 0.161 | -0.956 | -0.008 | 0.01 | 0.052 | 0.962 |

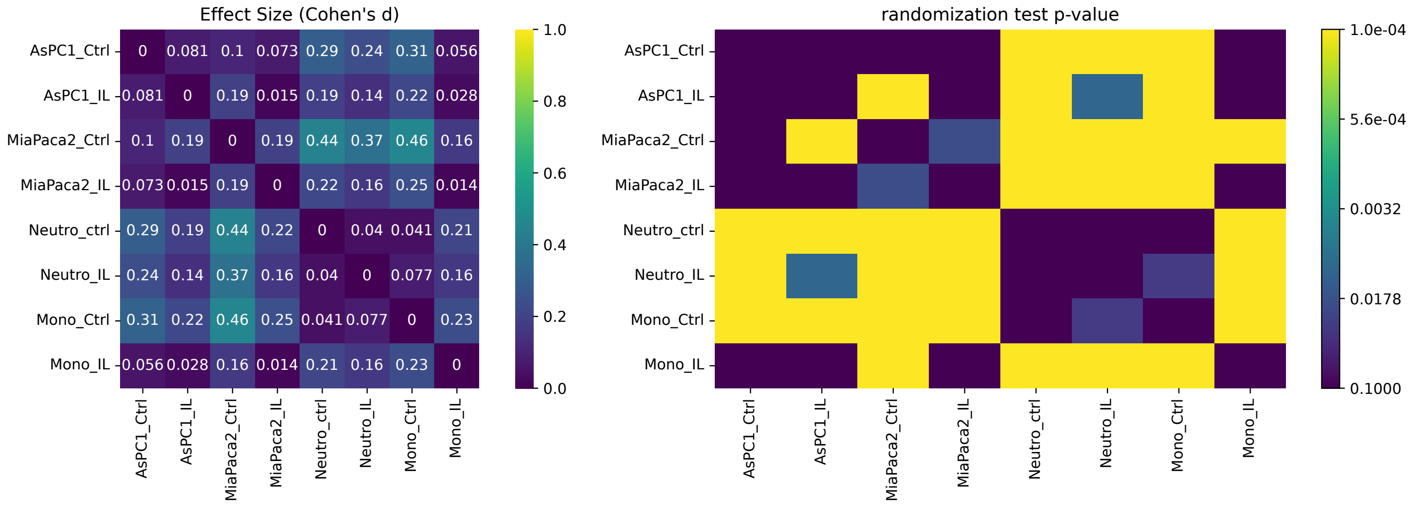


#### Fig3F

| Condition | count | mean | std | min | 25% | 50% | 75% | max |
|---|---|---|---|---|---|---|---|---|
| AsPC1 ctrl | 334.0 | 8.521 | 16.251 | 0.0 | 0.0 | 2.0 | 10.0 | 141.0 |
| MiaPaca2 ctrl | 395.0 | 22.152 | 35.776 | 0.0 | 2.0 | 7.0 | 26.5 | 253.0 |
| Mononuc. Ctrl | 359.0 | 3.443 | 8.924 | 0.0 | 0.0 | 1.0 | 3.5 | 102.0 |
| Neutro ctrl | 1710.0 | 11.991 | 15.856 | 0.0 | 2.0 | 6.0 | 15.0 | 136.0 |

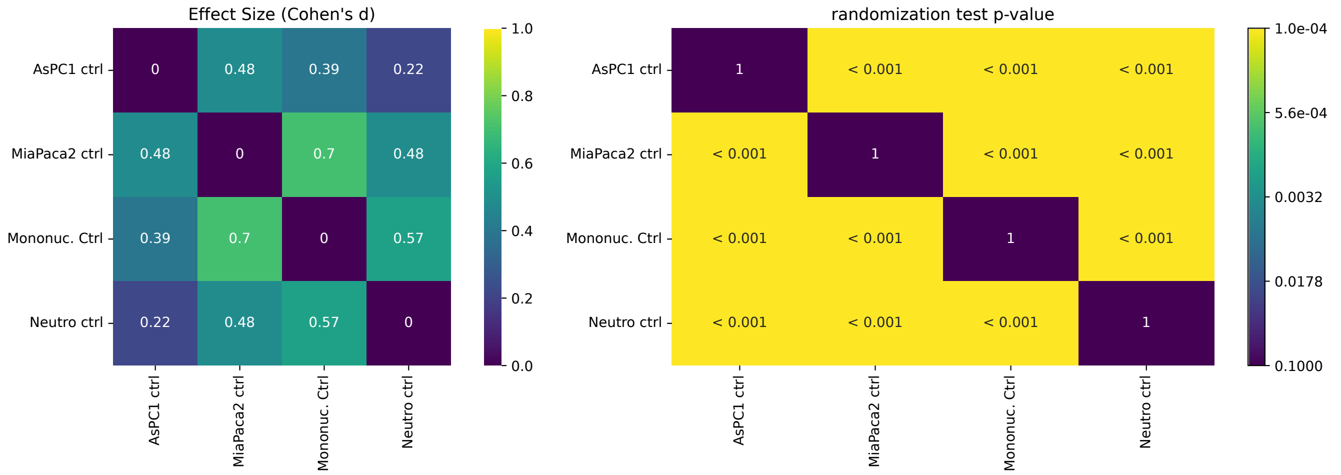

Fig4D

| Condition | count | mean | std | min | 25% | 50% | 75% | max |
|---|---|---|---|---|---|---|---|---|
| AsCP1 | 19.0 | 0.958 | 0.038 | 0.907 | 0.921 | 0.962 | 1.0 | 1.0 |
| BxPC3 | 26.0 | 0.985 | 0.031 | 0.895 | 1.0 | 1.0 | 1.0 | 1.0 |
| MiaPaca2 | 26.0 | 0.986 | 0.028 | 0.917 | 1.0 | 1.0 | 1.0 | 1.0 |
| PANC10.05 | 24.0 | 0.968 | 0.062 | 0.714 | 0.955 | 1.0 | 1.0 | 1.0 |
| PANC10.05.1 | 24.0 | 0.968 | 0.041 | 0.867 | 0.936 | 1.0 | 1.0 | 1.0 |
| SU86.86 | 22.0 | 0.973 | 0.056 | 0.8 | 1.0 | 1.0 | 1.0 | 1.0 |
| SW1990 | 24.0 | 0.977 | 0.029 | 0.92 | 0.956 | 1.0 | 1.0 | 1.0 |

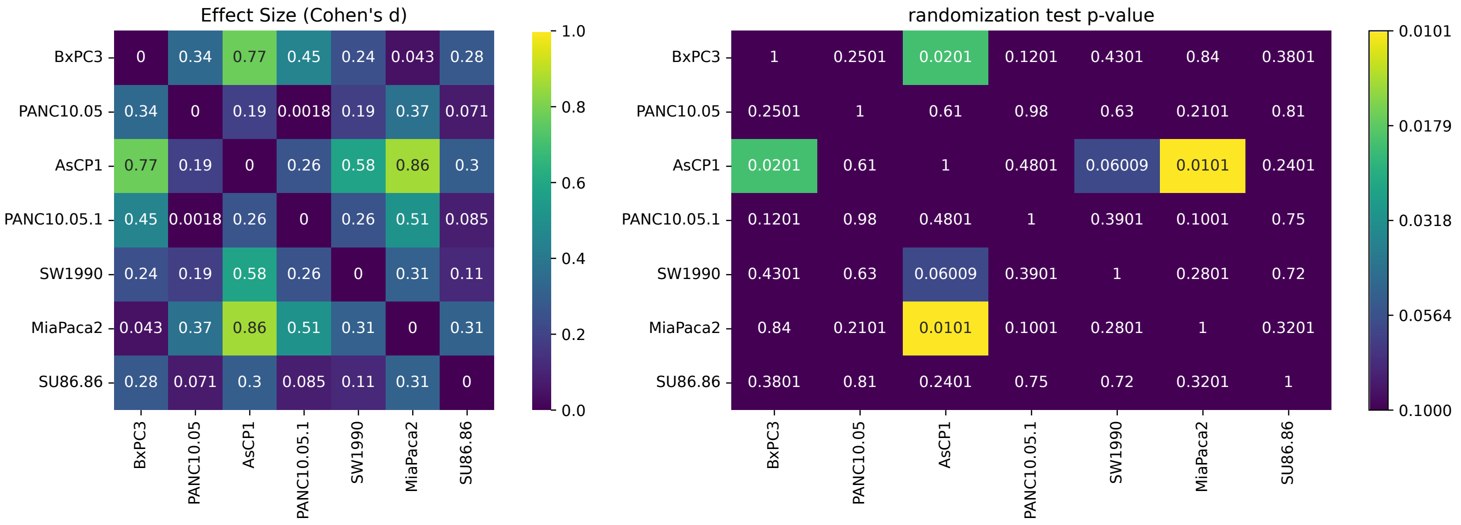

Fig5E

| Condition | count | mean | std | min | 25% | 50% | 75% | max |
| --- | --- | --- | --- | --- | --- | --- | --- | --- |
| cell body | 399.0 | 948.891 | 480.888 | 82.86 | 641.381 | 877.125 | 1159.725 | 2711.85 |
| junctions | 190.0 | 1218.288 | 497.12 | 338.299 | 831.775 | 1134.942 | 1522.008 | 2998.255 |

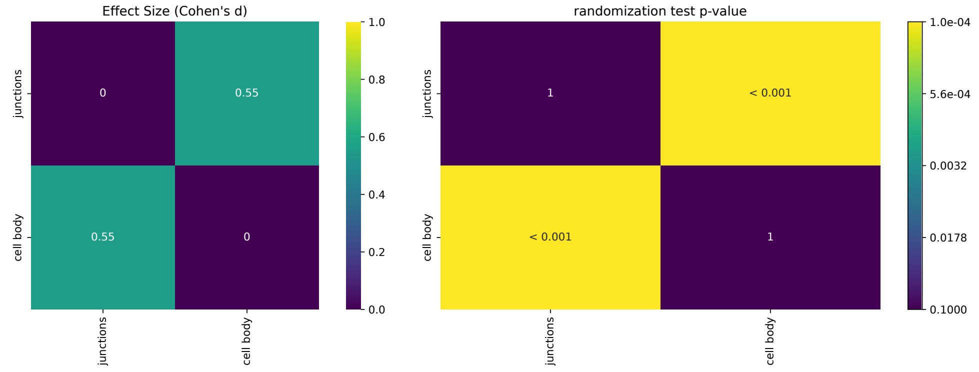


Fig5G2

| Condition | count | mean | std | min | 25% | 50% | 75% | max |
| --- | --- | --- | --- | --- | --- | --- | --- | --- |
| to junction | 1867.0 | 1.939 | 3.285 | 0.0 | 0.0 | 0.0 | 2.96 | 34.58 |
| to nuc | 1867.0 | 5.557 | 5.749 | 0.0 | 0.354 | 4.061 | 9.068 | 32.321 |

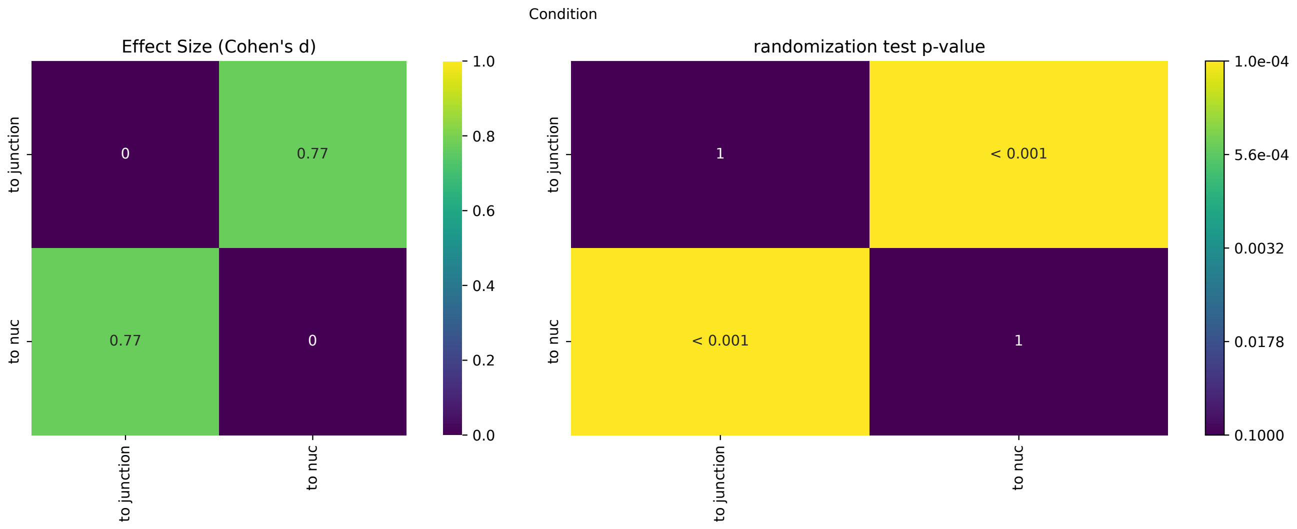

### Fig6D - MIA PaCa-2

| Condition | count | mean | std | min | 25% | 50% | 75% | max |
|---|---|---|---|---|---|---|---|---|
| CD44 expect. | 143.0 | 0.864 | 0.088 | 0.575 | 0.8 | 0.896 | 0.934 | 0.977 |
| CD44 obs. | 143.0 | 0.972 | 0.04 | 0.833 | 0.947 | 1.0 | 1.0 | 1.0 |
| FN expect. | 135.0 | 0.153 | 0.043 | 0.054 | 0.126 | 0.154 | 0.178 | 0.261 |
| FN obs | 135.0 | 0.185 | 0.099 | 0.03 | 0.115 | 0.16 | 0.231 | 0.522 |
| ICAM1 expect. | 105.0 | 0.342 | 0.181 | 0.071 | 0.173 | 0.322 | 0.492 | 0.815 |
| ICAM1 obs | 105.0 | 0.19 | 0.149 | 0.0 | 0.107 | 0.155 | 0.243 | 1.0 |
| ICAM2 expect. | 113.0 | 1.0 | 0.0 | 1.0 | 1.0 | 1.0 | 1.0 | 1.0 |
| ICAM2 obs | 113.0 | 1.0 | 0.0 | 1.0 | 1.0 | 1.0 | 1.0 | 1.0 |

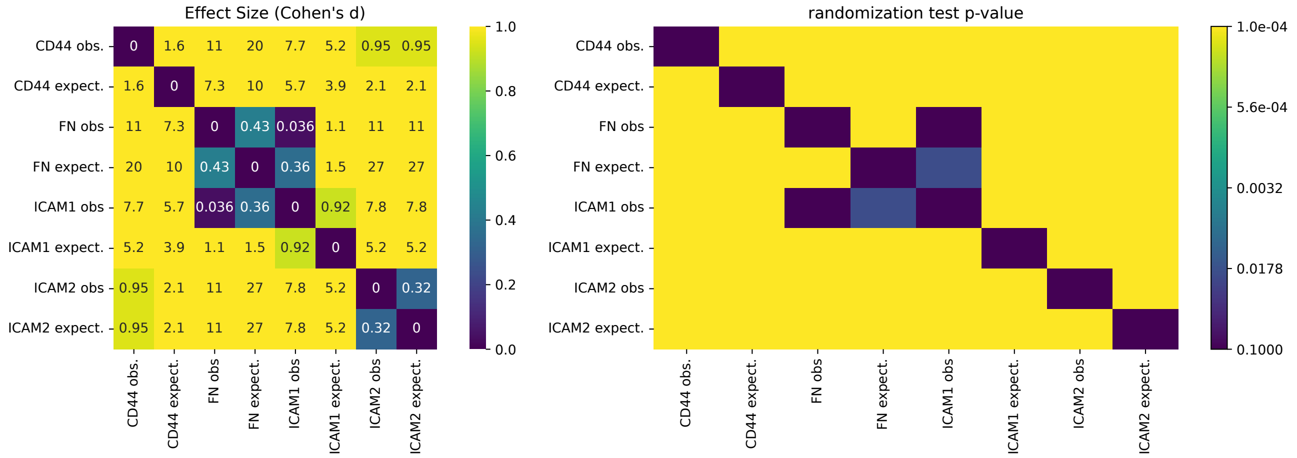


### Fig6D - AsPC-1

| Condition | count | mean | std | min | 25% | 50% | 75% | max |
|---|---|---|---|---|---|---|---|---|
| CD44 expect. | 143.0 | 0.872 | 0.07 | 0.575 | 0.832 | 0.868 | 0.928 | 0.989 |
| CD44 obs. | 143.0 | 0.963 | 0.061 | 0.667 | 0.946 | 1.0 | 1.0 | 1.0 |
| FN expect. | 135.0 | 0.135 | 0.039 | 0.054 | 0.108 | 0.129 | 0.16 | 0.286 |
| FN obs | 135.0 | 0.236 | 0.152 | 0.0 | 0.13 | 0.2 | 0.296 | 0.765 |
| ICAM1 expect. | 105.0 | 0.17 | 0.059 | 0.044 | 0.123 | 0.166 | 0.211 | 0.307 |
| ICAM1 obs | 105.0 | 0.174 | 0.145 | 0.0 | 0.091 | 0.161 | 0.243 | 1.0 |
| ICAM2 expect. | 113.0 | 1.0 | 0.001 | 0.997 | 1.0 | 1.0 | 1.0 | 1.0 |
| ICAM2 obs | 113.0 | 1.0 | 0.0 | 1.0 | 1.0 | 1.0 | 1.0 | 1.0 |

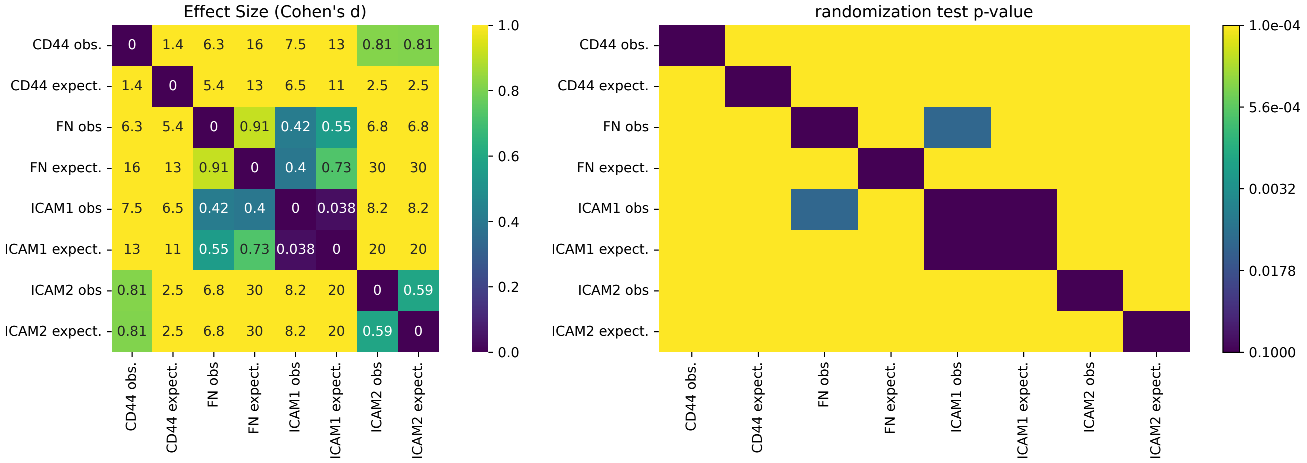

Fig7C

| Condition | count | mean | std | min | 25% | 50% | 75% | max |
|---|---|---|---|---|---|---|---|---|
| As 200 si1 | 3.0 | 0.018 | 0.03 | -0.0 | 0.0 | 0.001 | 0.027 | 0.053 |
| As 200 si2 | 3.0 | 0.01 | 0.014 | -0.006 | 0.004 | 0.014 | 0.018 | 0.022 |
| As 200 siCtrl | 3.0 | 0.621 | 0.135 | 0.497 | 0.549 | 0.6 | 0.682 | 0.764 |
| Mia 200 si1 | 3.0 | -0.006 | 0.004 | -0.011 | -0.008 | -0.004 | -0.004 | -0.003 |
| Mia 200 si2 | 3.0 | 0.006 | 0.011 | -0.005 | 0.002 | 0.009 | 0.012 | 0.016 |
| Mia 200 siCtrl | 3.0 | 2.087 | 0.559 | 1.684 | 1.768 | 1.853 | 2.289 | 2.725 |

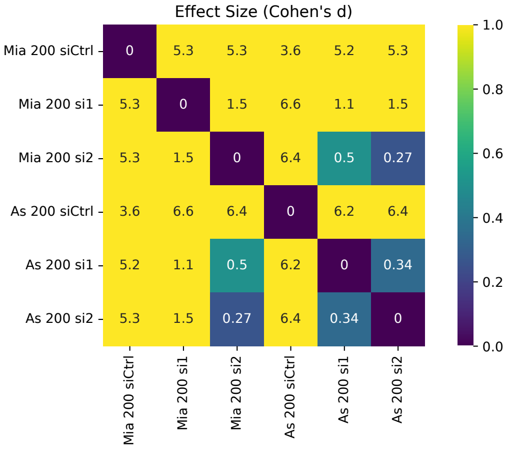


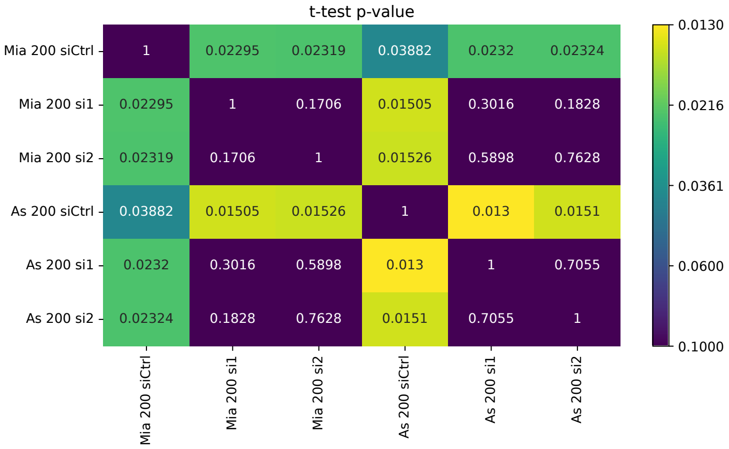


Fig7F

| Condition | count | mean | std | min | 25% | 50% | 75% | max |
|---|---|---|---|---|---|---|---|---|
| AsPc1_200_HUblock | 3.0 | 1.112 | 0.206 | 0.912 | 1.006 | 1.1 | 1.212 | 1.323 |
| AsPc1_200_TCblock | 3.0 | 0.025 | 0.035 | -0.007 | 0.006 | 0.02 | 0.041 | 0.062 |
| AsPc1_200_blockboth | 2.0 | -0.003 | 0.01 | -0.009 | -0.006 | -0.003 | 0.001 | 0.004 |
| AsPc1_200_ctrlblock | 2.0 | 0.607 | 0.215 | 0.455 | 0.531 | 0.607 | 0.683 | 0.759 |
| Mia_200_HUblock | 4.0 | 1.658 | 1.165 | 0.825 | 0.898 | 1.234 | 1.994 | 3.338 |
| Mia_200_TCblock | 4.0 | 0.036 | 0.026 | 0.003 | 0.022 | 0.041 | 0.056 | 0.061 |
| Mia_200_blockboth | 4.0 | 0.017 | 0.021 | -0.0 | 0.005 | 0.01 | 0.022 | 0.048 |
| Mia_200_ctrlblock | 5.0 | 1.282 | 0.646 | 0.283 | 1.229 | 1.32 | 1.516 | 2.063 |

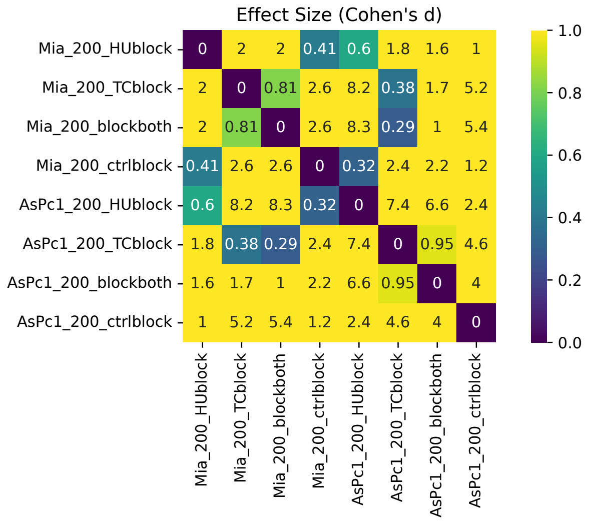


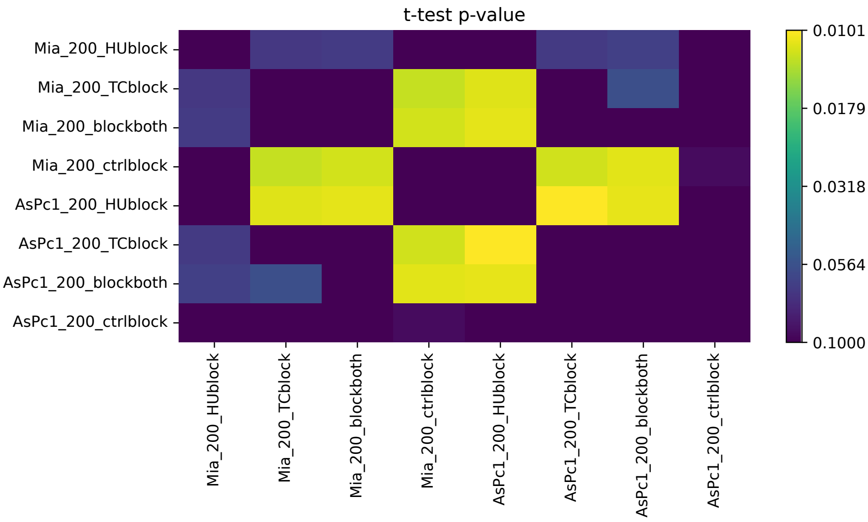

#### Fig7G

| Condition | count | mean | std | min | 25% | 50% | 75% | max |
|---|---|---|---|---|---|---|---|---|
| As CD44 2 | 12.0 | 119.627 | 57.985 | 37.788 | 90.062 | 122.739 | 147.073 | 244.346 |
| As Ctrl | 12.0 | 53.072 | 32.65 | 22.916 | 37.477 | 42.761 | 58.671 | 140.363 |
| As siCD44 1 | 12.0 | 138.203 | 99.062 | -50.0 | 66.672 | 148.596 | 213.27 | 271.691 |
| Mia Ctrl | 12.0 | 30.057 | 22.139 | 10.67 | 13.159 | 21.513 | 38.095 | 77.139 |
| Mia siCD44 1 | 12.0 | 148.547 | 77.603 | 14.919 | 100.661 | 144.835 | 196.574 | 291.173 |
| Mia siCD44 2 | 12.0 | 111.535 | 71.089 | 23.681 | 78.854 | 109.534 | 133.61 | 282.099 |

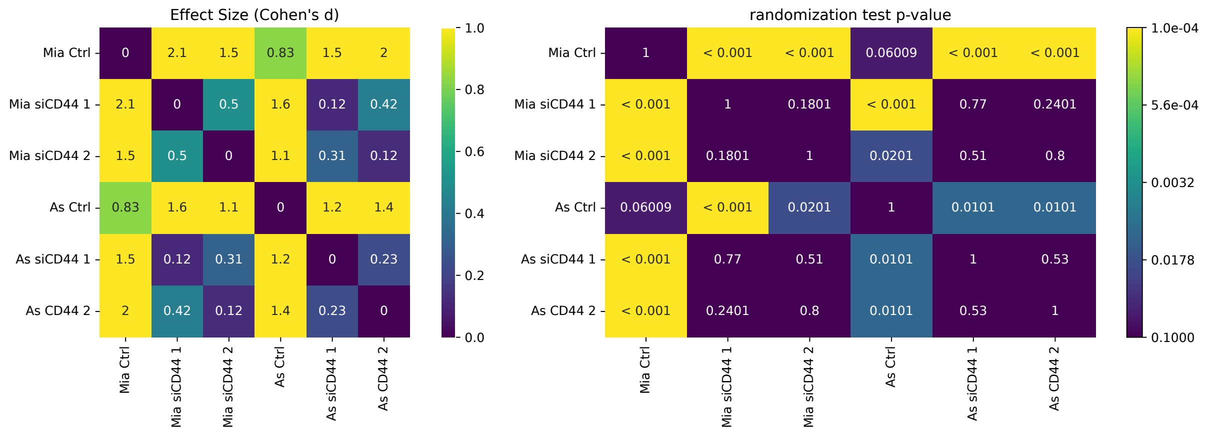


#### Fig7H

| Condition | count | mean | std | min | 25% | 50% | 75% | max |
|---|---|---|---|---|---|---|---|---|
| As CD44 2 | 79.0 | 120.361 | 189.809 | 0.0 | 7.764 | 38.121 | 133.078 | 1099.91 |
| As Ctrl | 838.0 | 184.71 | 166.411 | 0.0 | 52.281 | 136.616 | 281.322 | 1087.24 |
| As siCD44 1 | 88.0 | 126.488 | 145.42 | 0.556 | 16.018 | 68.668 | 167.461 | 500.726 |
| Mia Ctrl | 2172.0 | 179.863 | 155.522 | 0.138 | 59.259 | 130.928 | 269.529 | 1023.384 |
| Mia siCD44 1 | 35.0 | 90.018 | 138.32 | 0.0 | 4.505 | 30.417 | 96.632 | 573.91 |
| Mia siCD44 2 | 98.0 | 135.73 | 173.008 | 0.0 | 19.646 | 74.844 | 170.194 | 841.671 |

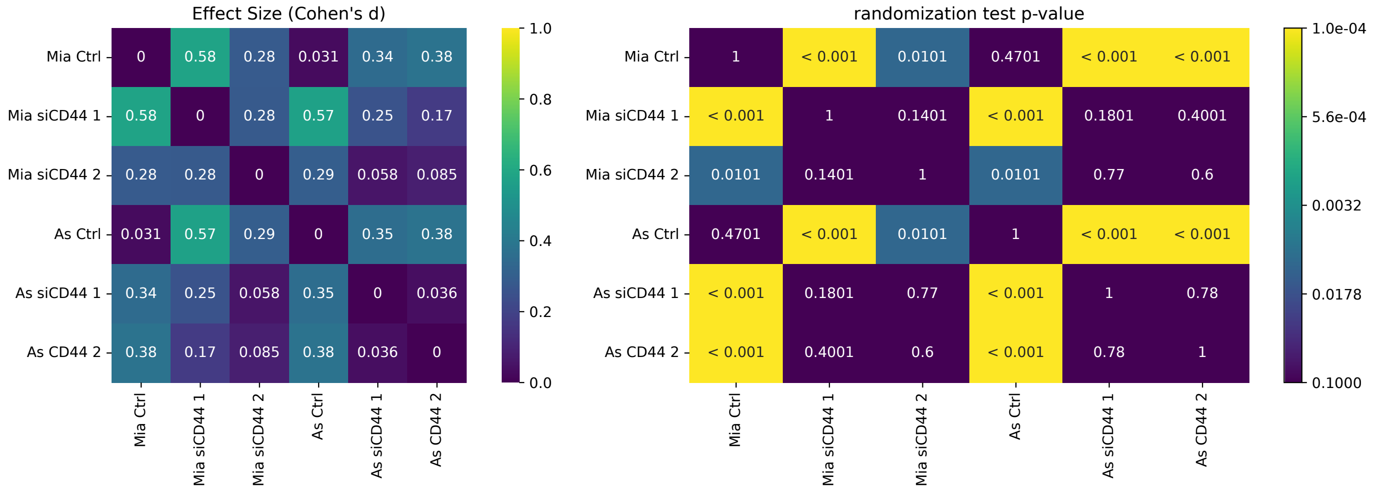

### Fig7I

| Condition | count | mean | std | min | 25% | 50% | 75% | max |
|---|---|---|---|---|---|---|---|---|
| As CD44 2 | 79.0 | 26.253 | 40.411 | 0.0 | 2.0 | 9.0 | 29.0 | 210.0 |
| As Ctrl | 838.0 | 40.599 | 34.719 | 0.0 | 12.0 | 31.0 | 62.0 | 173.0 |
| As siCD44 1 | 88.0 | 29.307 | 34.229 | 0.0 | 4.75 | 16.5 | 37.5 | 123.0 |
| Mia Ctrl | 2172.0 | 39.629 | 32.955 | 0.0 | 14.0 | 30.0 | 60.0 | 195.0 |
| Mia siCD44 1 | 35.0 | 22.143 | 37.447 | 0.0 | 1.5 | 5.0 | 24.0 | 152.0 |
| Mia siCD44 2 | 98.0 | 26.888 | 31.812 | 0.0 | 4.0 | 14.0 | 38.0 | 145.0 |

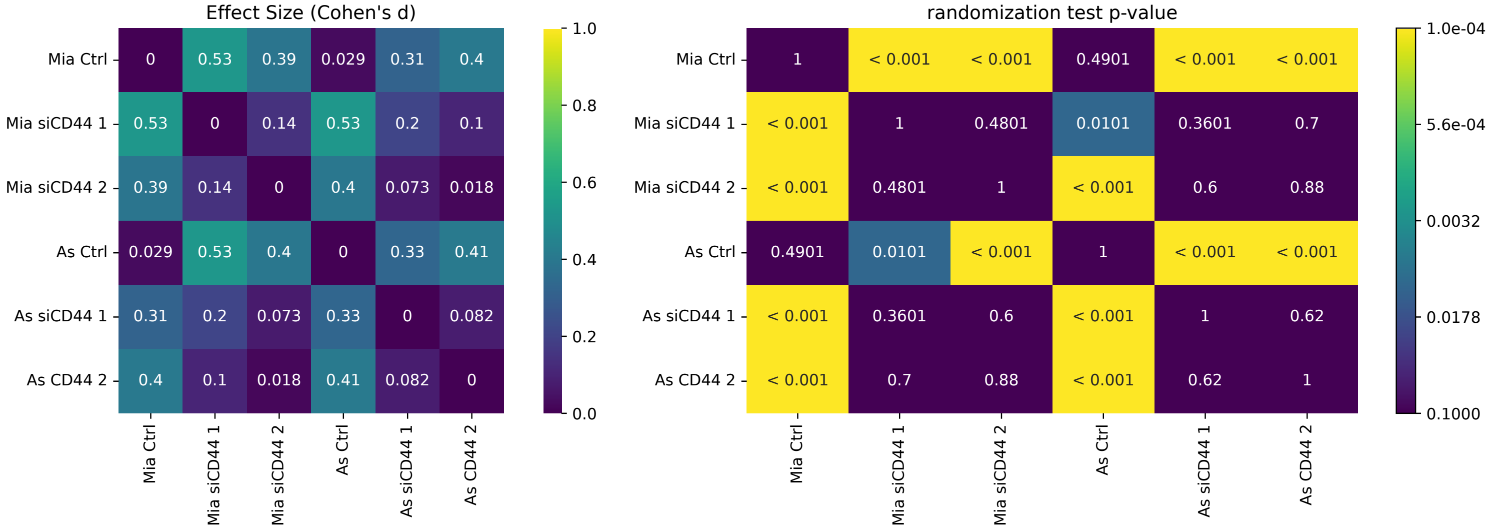


### Fig7J

| Condition | count | mean | std | min | 25% | 50% | 75% | max |
|---|---|---|---|---|---|---|---|---|
| Bothblock | 8.0 | 144.735 | 62.447 | 48.96 | 122.92 | 150.452 | 177.271 | 242.517 |
| Bothblock Mia | 16.0 | 156.435 | 169.38 | 17.933 | 68.304 | 94.999 | 147.823 | 628.896 |
| Ctrlblock | 8.0 | 58.471 | 30.202 | 35.75 | 39.609 | 45.267 | 66.149 | 124.166 |
| Ctrlblock Mia | 20.0 | 52.537 | 24.436 | 25.46 | 35.783 | 48.525 | 62.882 | 117.376 |
| HU block | 12.0 | 57.141 | 34.46 | 23.852 | 38.596 | 43.768 | 60.844 | 139.168 |
| HU block Mia | 16.0 | 62.53 | 39.503 | 27.489 | 31.887 | 46.839 | 79.724 | 148.968 |
| TC block | 12.0 | 118.615 | 60.12 | 30.978 | 78.184 | 121.923 | 155.304 | 233.0 |
| TC block Mia | 16.0 | 108.221 | 60.094 | 14.839 | 55.78 | 119.032 | 147.303 | 232.326 |

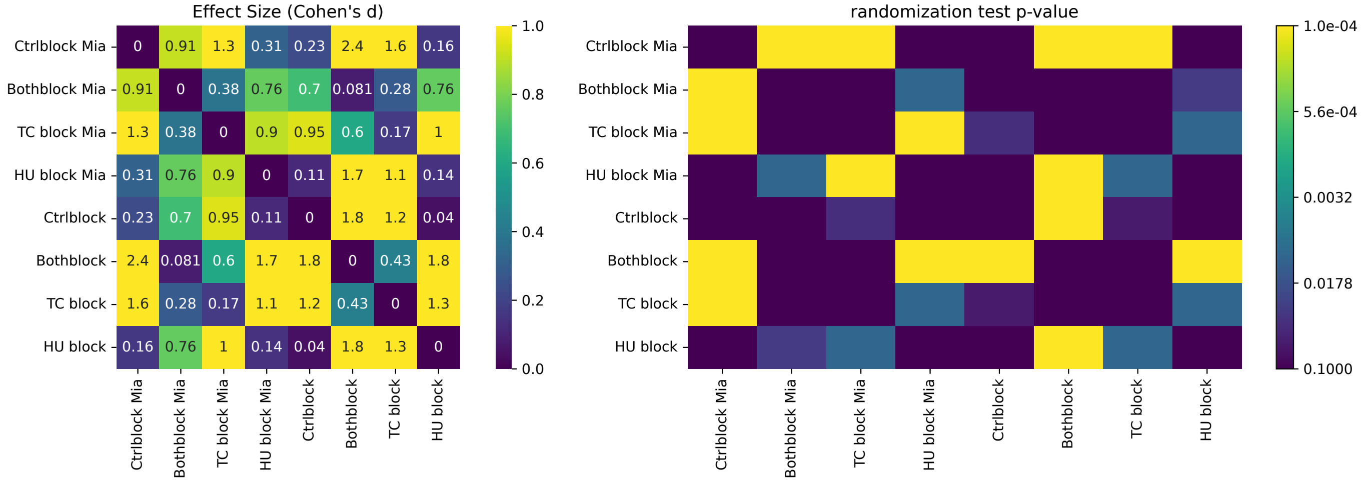

#### Fig7K

| Condition | count | mean | std | min | 25% | 50% | 75% | max |
|---|---|---|---|---|---|---|---|---|
| Bothblock As | 35.0 | 86.618 | 134.821 | 0.98 | 10.318 | 36.729 | 91.239 | 667.249 |
| Bothblock Mia | 227.0 | 112.14 | 136.938 | 0.0 | 20.009 | 61.29 | 160.377 | 954.225 |
| Ctrlblock As | 673.0 | 159.525 | 157.75 | 0.072 | 41.973 | 106.053 | 235.992 | 963.331 |
| Ctrlblock Mia | 1721.0 | 179.763 | 159.206 | 0.137 | 59.704 | 140.583 | 259.506 | 1627.397 |
| HU block As | 993.0 | 186.249 | 160.931 | 0.0 | 56.757 | 143.593 | 279.998 | 1226.981 |
| HU block Mia | 1525.0 | 182.057 | 151.37 | 0.0 | 61.511 | 147.567 | 272.943 | 1211.935 |
| TC block As | 101.0 | 151.295 | 262.536 | 0.0 | 12.92 | 50.326 | 174.886 | 1583.529 |
| TC block Mia | 161.0 | 87.646 | 106.02 | 0.0 | 15.369 | 44.781 | 122.087 | 605.513 |

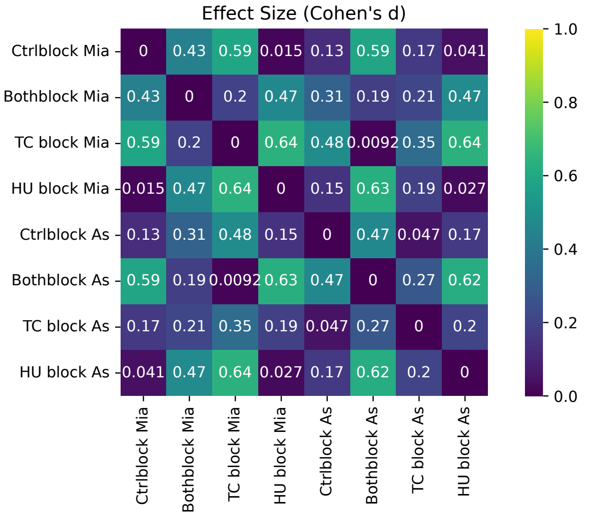


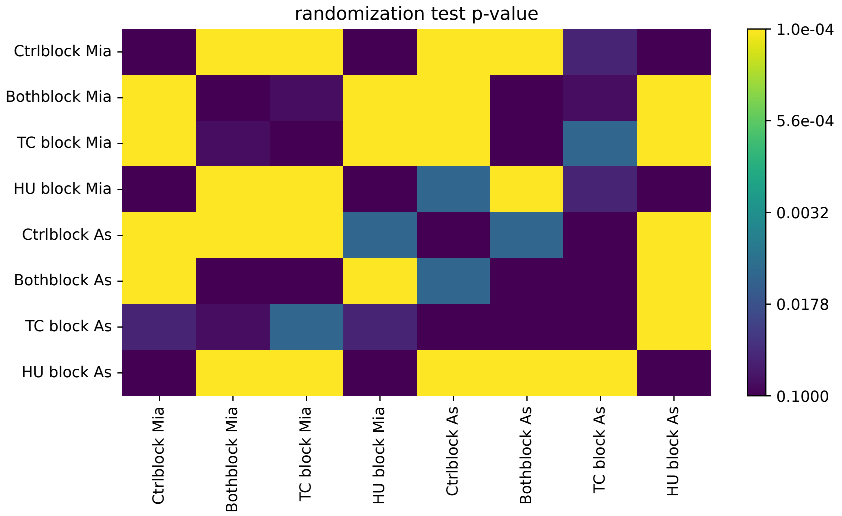


#### Fig7L

| Condition | count | mean | std | min | 25% | 50% | 75% | max |
|---|---|---|---|---|---|---|---|---|
| Bothblock As | 35.0 | 20.571 | 38.375 | 0.0 | 3.0 | 8.0 | 21.0 | 212.0 |
| Ctrlblock | 1721.0 | 39.792 | 32.042 | 0.0 | 14.0 | 32.0 | 57.0 | 178.0 |
| Ctrlblock As | 673.0 | 34.978 | 32.351 | 0.0 | 10.0 | 25.0 | 54.0 | 155.0 |
| ECblock | 1525.0 | 39.361 | 30.188 | 0.0 | 14.0 | 33.0 | 59.0 | 169.0 |
| HU block As | 993.0 | 40.836 | 33.51 | 0.0 | 13.0 | 33.0 | 61.0 | 189.0 |
| TCblock | 161.0 | 20.609 | 25.623 | 0.0 | 4.0 | 11.0 | 26.0 | 173.0 |
| TC block As | 101.0 | 28.644 | 38.019 | 0.0 | 3.0 | 11.0 | 41.0 | 149.0 |
| bothblock | 227.0 | 22.529 | 23.509 | 0.0 | 5.0 | 14.0 | 33.0 | 110.0 |

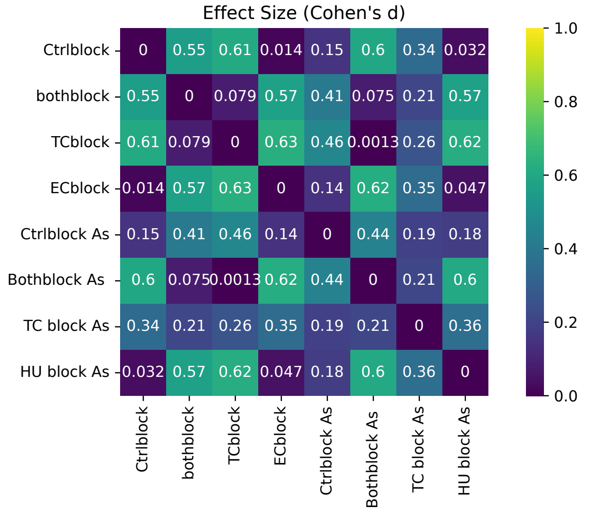


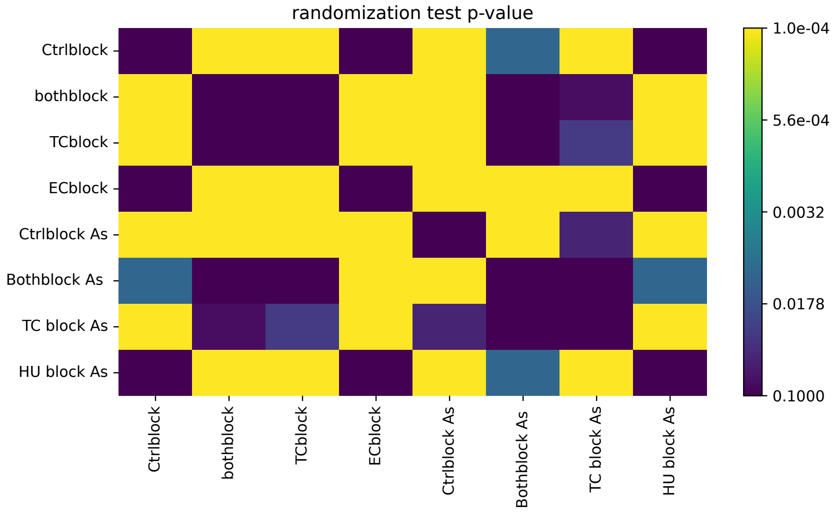

#### Fig8B-AsPC1

| Condition | count | mean | std | min | 25% | 50% | 75% | max |
|---|---|---|---|---|---|---|---|---|
| 120U | 1397.0 | 0.767 | 0.658 | 0.0 | 0.373 | 0.529 | 0.884 | 5.04 |
| Untreated | 1542.0 | 1.0 | 0.655 | 0.0 | 0.552 | 0.846 | 1.267 | 5.567 |

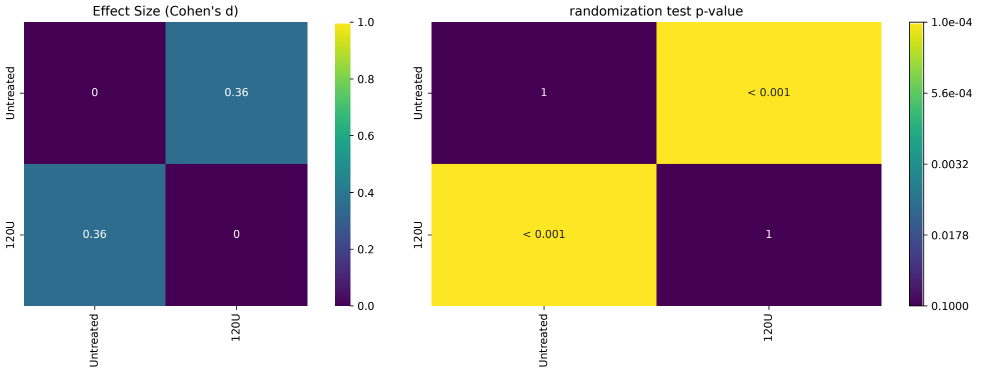


#### Fig8B-MiaPaCa2

| Condition | count | mean | std | min | 25% | 50% | 75% | max |
|---|---|---|---|---|---|---|---|---|
| 120U | 601.0 | 0.737 | 0.69 | 0.0 | 0.308 | 0.468 | 0.883 | 5.324 |
| Untreated | 959.0 | 1.0 | 0.926 | 0.0 | 0.383 | 0.73 | 1.351 | 8.63 |

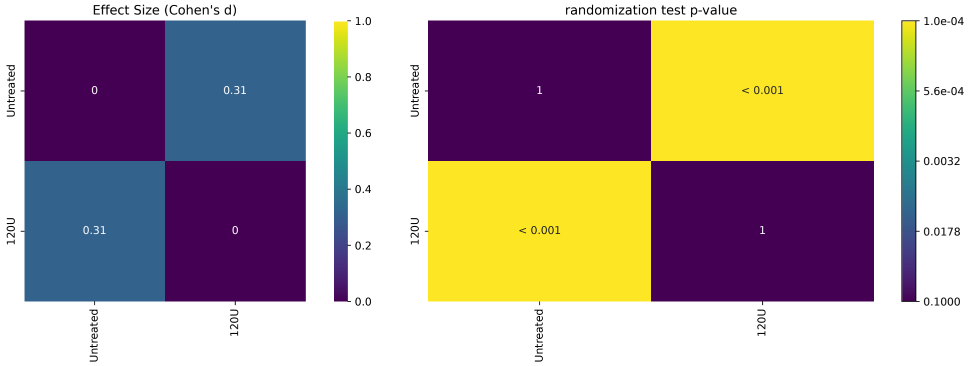

Fig8G

| Condition | count | mean | std | min | 25% | 50% | 75% | max |
|---|---|---|---|---|---|---|---|---|
| Ctrl dig As | 754.0 | 169.043 | 153.843 | 0.0 | 48.487 | 128.285 | 243.606 | 1201.998 |
| Ctrl dig Mia | 1055.0 | 192.58 | 165.878 | 0.137 | 66.733 | 153.644 | 285.173 | 1333.766 |
| EC dig As | 41.0 | 121.082 | 118.908 | 0.844 | 33.63 | 100.318 | 159.273 | 552.231 |
| EC dig Mia | 45.0 | 86.669 | 189.905 | 0.643 | 16.04 | 32.545 | 61.928 | 1188.622 |
| TC dig As | 20.0 | 85.901 | 85.78 | 0.0 | 17.246 | 48.062 | 143.873 | 265.651 |
| TC dig Mia | 77.0 | 83.662 | 109.024 | 0.0 | 6.624 | 41.439 | 113.179 | 452.494 |

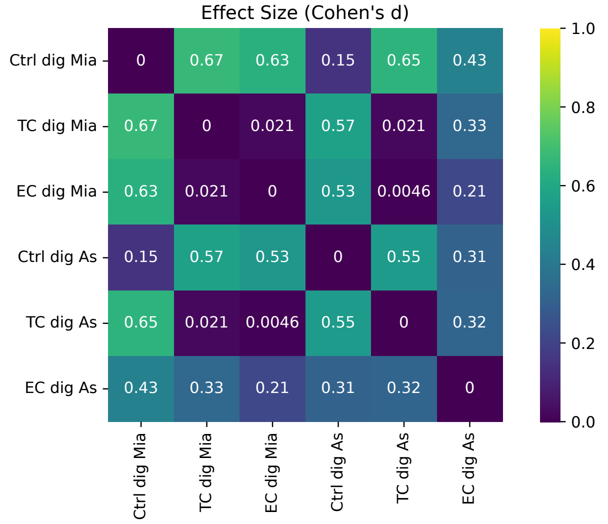


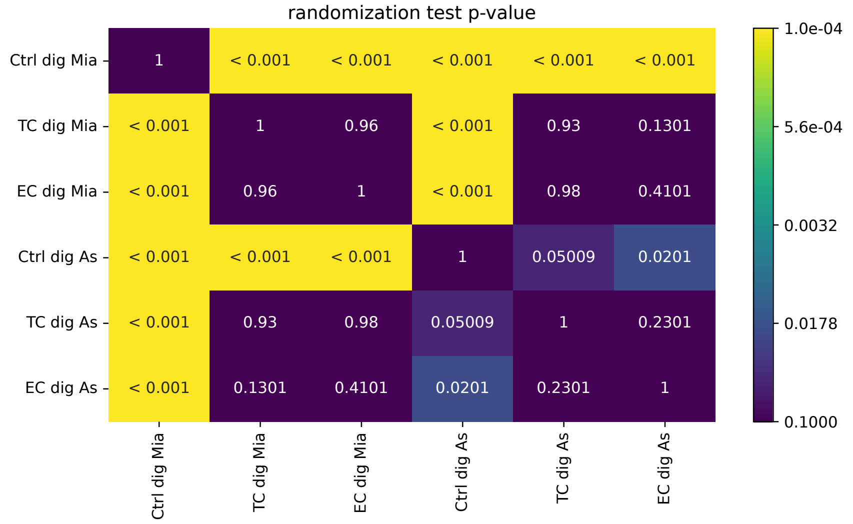


Fig8H

| Condition | count | mean | std | min | 25% | 50% | 75% | max |
|---|---|---|---|---|---|---|---|---|
| Ctrl dig As | 754.0 | 40.219 | 34.759 | 0.0 | 12.0 | 32.0 | 58.0 | 190.0 |
| Ctrl dig Mia | 1055.0 | 41.898 | 31.836 | 0.0 | 16.0 | 36.0 | 63.0 | 195.0 |
| EC dig As | 41.0 | 26.634 | 26.189 | 0.0 | 7.0 | 21.0 | 41.0 | 132.0 |
| EC dig Mia | 45.0 | 15.089 | 20.938 | 0.0 | 3.0 | 9.0 | 15.0 | 94.0 |
| TC dig As | 20.0 | 20.4 | 19.019 | 0.0 | 5.0 | 14.0 | 29.5 | 60.0 |
| TC dig Mia | 77.0 | 18.896 | 26.297 | 0.0 | 2.0 | 8.0 | 26.0 | 127.0 |

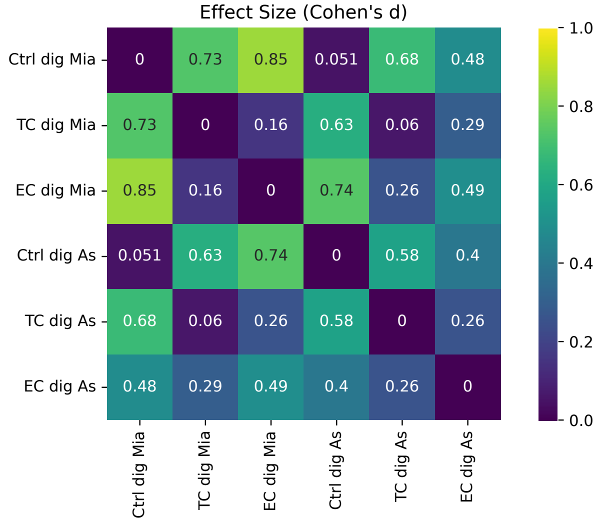


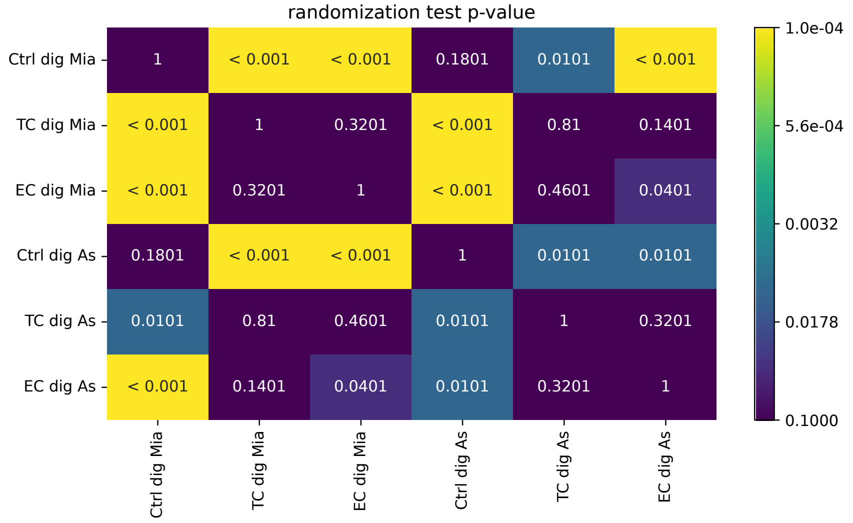

#### Fig8E

| Condition | count | mean | std | min | 25% | 50% | 75% | max |
|---|---|---|---|---|---|---|---|---|
| AsPc1_200_ECdig | 3.0 | 0.002 | 0.005 | -0.001 | -0.001 | -0.001 | 0.004 | 0.009 |
| AsPc1_200_TCdig | 2.0 | -0.025 | 0.025 | -0.042 | -0.034 | -0.025 | -0.016 | -0.007 |
| AsPc1_200_ctrldig | 3.0 | 0.784 | 0.82 | 0.155 | 0.321 | 0.486 | 1.099 | 1.711 |
| Mia_200_ECdig | 3.0 | 0.011 | 0.008 | 0.003 | 0.007 | 0.011 | 0.015 | 0.019 |
| Mia_200_TCdig | 3.0 | -0.014 | 0.041 | -0.059 | -0.033 | -0.007 | 0.008 | 0.022 |
| Mia_200_ctrldig | 3.0 | 1.122 | 0.281 | 0.801 | 1.023 | 1.244 | 1.282 | 1.32 |

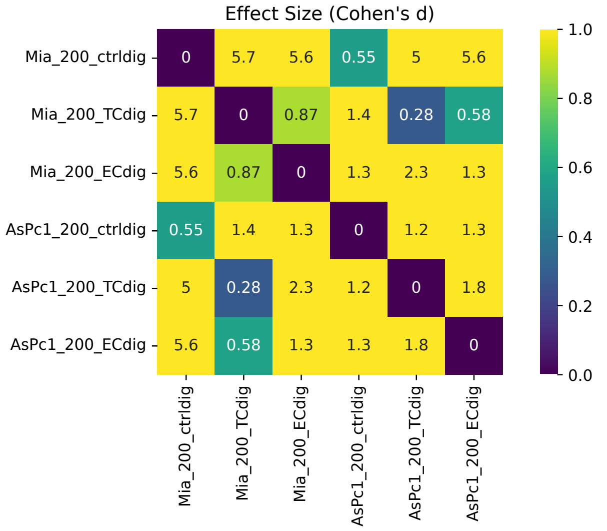


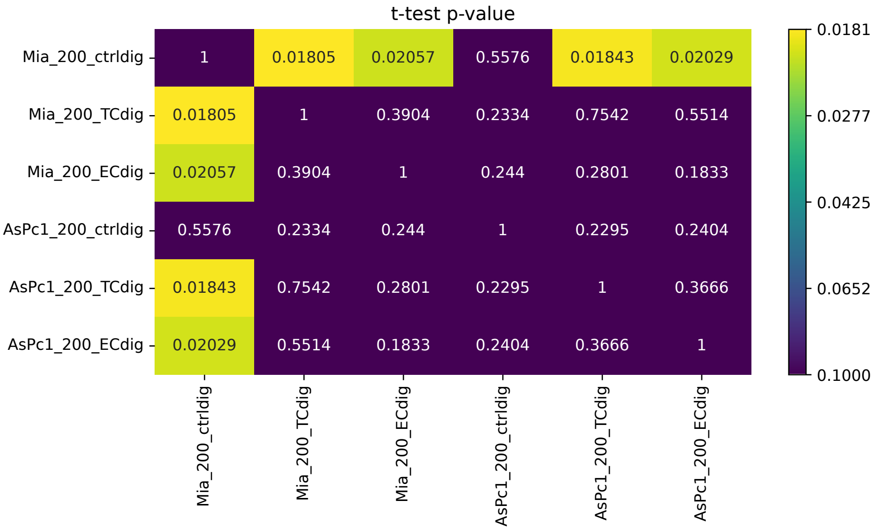


#### Fig8F

| Condition | count | mean | std | min | 25% | 50% | 75% | max |
|---|---|---|---|---|---|---|---|---|
| Ctrl dig As | 12.0 | 80.785 | 45.373 | 31.067 | 48.088 | 72.334 | 100.881 | 162.895 |
| Ctrl dig Mia | 12.0 | 58.399 | 44.301 | 22.246 | 26.1 | 40.537 | 66.314 | 153.335 |
| EC dig As | 12.0 | 142.386 | 139.139 | -50.0 | 60.306 | 125.863 | 252.747 | 412.244 |
| EC dig Mia | 12.0 | 150.125 | 65.666 | 41.179 | 115.104 | 151.634 | 207.831 | 239.902 |
| TC dig As | 8.0 | 146.794 | 100.836 | 28.589 | 67.498 | 129.27 | 224.072 | 294.995 |
| TC dig Mia | 12.0 | 116.091 | 61.668 | 32.556 | 67.5 | 110.277 | 171.967 | 206.511 |

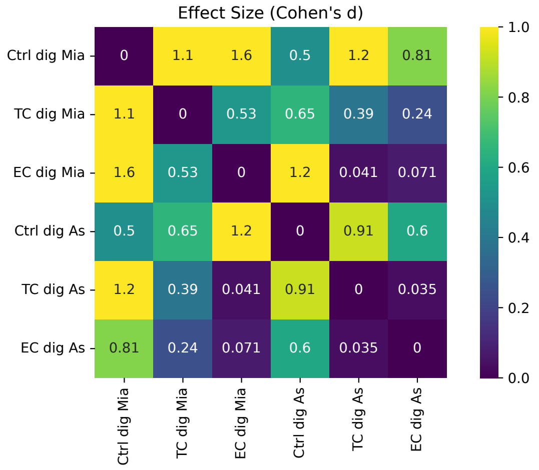


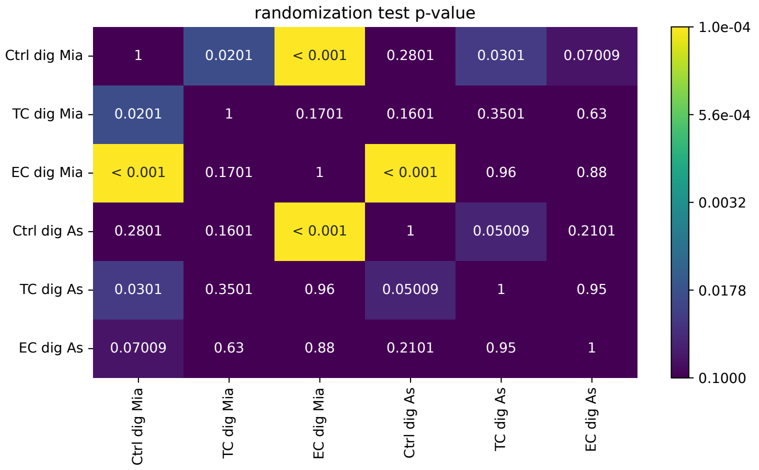
