## Supplementary material for "Fast label-free live imaging reveals key roles of flow dynamics and CD44-HA interaction in cancer cell arrest on endothelial monolayers": SI figures Statistical summaries

### Figure S1 to figure S8, Statistical summaries

FigS2E

| Condition | count | mean | std | min | 25% | 50% | 75% | max |
|---|---|---|---|---|---|---|---|---|
| AsPC1 | 18.0 | 1067.222 | 690.892 | 142.0 | 270.25 | 1119.5 | 1709.75 | 2009.0 |
| BxPC3 | 15.0 | 175.067 | 55.214 | 65.0 | 130.5 | 187.0 | 202.0 | 269.0 |
| MiaPaca2 | 23.0 | 500.043 | 342.065 | 115.0 | 297.0 | 415.0 | 530.0 | 1518.0 |
| PANC1 | 18.0 | 197.833 | 68.479 | 108.0 | 159.75 | 179.5 | 228.5 | 401.0 |
| PANC10.05 | 27.0 | 174.148 | 67.424 | 36.0 | 134.0 | 172.0 | 199.0 | 335.0 |
| SU8686 | 10.0 | 110.0 | 62.279 | 16.0 | 75.5 | 96.5 | 125.0 | 223.0 |
| SW1990 | 16.0 | 576.75 | 358.421 | 69.0 | 265.75 | 632.5 | 858.25 | 1244.0 |

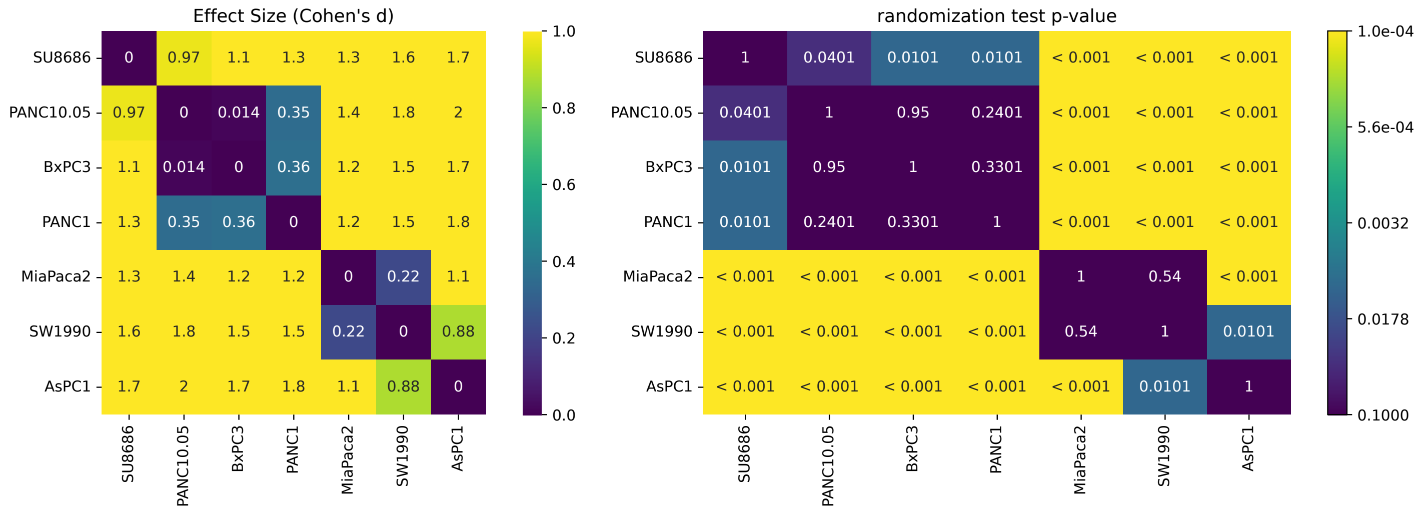


FigS2F

| Condition | count | mean | std | min | 25% | 50% | 75% | max |
|---|---|---|---|---|---|---|---|---|
| AsPC1 | 39.0 | 11.737 | 2.189 | 8.92 | 10.2 | 10.949 | 13.322 | 18.248 |
| BxPC3 | 40.0 | 13.736 | 2.146 | 9.372 | 12.539 | 13.689 | 15.021 | 19.544 |
| MiaPaca2 | 40.0 | 13.775 | 3.106 | 8.859 | 11.738 | 13.4 | 15.252 | 23.736 |
| PANC1 | 40.0 | 19.387 | 4.639 | 13.159 | 16.077 | 18.444 | 21.279 | 31.89 |
| PANC10.05 | 40.0 | 16.036 | 2.137 | 11.588 | 14.497 | 15.846 | 17.41 | 22.329 |
| SU8686 | 40.0 | 18.875 | 2.657 | 13.813 | 17.11 | 18.405 | 21.041 | 23.91 |
| SW1990 | 40.0 | 17.598 | 4.389 | 11.031 | 14.419 | 16.272 | 19.38 | 27.918 |

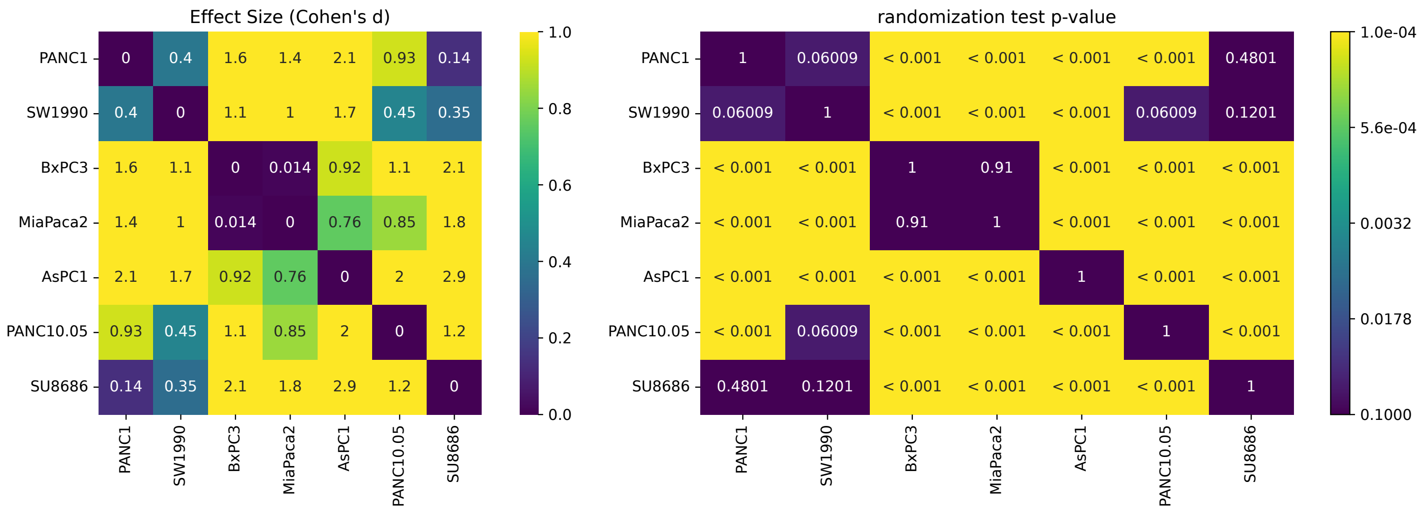

### FigS3B_MaxSpeed

| Condition | count | mean | std | min | 25% | 50% | 75% | max |
|---|---|---|---|---|---|---|---|---|
| As_100 | 7329.0 | 103.041 | 85.074 | 4.109 | 23.896 | 94.02 | 140.883 | 374.481 |
| As_200 | 7329.0 | 214.44 | 94.723 | 5.624 | 171.253 | 219.24 | 284.033 | 374.934 |
| As_300 | 6854.0 | 266.752 | 83.281 | 4.908 | 230.563 | 284.567 | 326.773 | 374.958 |
| As_wash | 5325.0 | 121.798 | 126.147 | 4.084 | 24.422 | 47.188 | 255.839 | 374.947 |
| Mia_100 | 3010.0 | 25.336 | 41.12 | 2.995 | 9.534 | 14.227 | 20.47 | 372.208 |
| Mia_200 | 3232.0 | 55.395 | 72.676 | 3.503 | 15.382 | 25.31 | 56.799 | 374.915 |
| Mia_300 | 3232.0 | 83.633 | 92.799 | 4.781 | 20.422 | 42.037 | 105.002 | 374.943 |
| Mia_wash | 4732.0 | 33.403 | 59.412 | 4.136 | 10.477 | 15.681 | 24.796 | 374.896 |
| P10_100 | 8226.0 | 168.412 | 75.728 | 11.621 | 114.457 | 144.804 | 212.54 | 374.663 |
| P10_200 | 8226.0 | 227.344 | 71.182 | 15.117 | 185.833 | 229.465 | 275.446 | 374.943 |
| P10_300 | 6857.0 | 264.431 | 64.224 | 6.738 | 217.593 | 265.122 | 317.387 | 374.943 |
| P10_wash | 5935.0 | 287.22 | 52.352 | 15.542 | 245.44 | 289.522 | 331.183 | 374.963 |

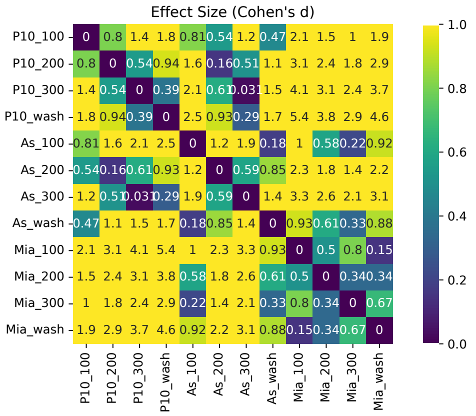


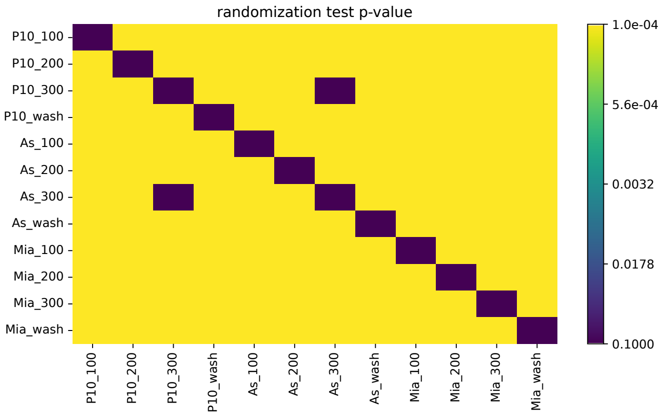


### FigS3B_MeanSpeed

| Condition | count | mean | std | min | 25% | 50% | 75% | max |
|---|---|---|---|---|---|---|---|---|
| As_100 | 7329.0 | 33.941 | 28.032 | 1.163 | 3.676 | 36.192 | 55.64 | 194.819 |
| As_200 | 7329.0 | 100.258 | 48.992 | 1.723 | 83.102 | 110.341 | 133.102 | 250.796 |
| As_300 | 6854.0 | 134.802 | 60.735 | 1.388 | 93.332 | 135.378 | 179.239 | 311.593 |
| As_wash | 5325.0 | 40.628 | 70.24 | 1.257 | 3.858 | 5.431 | 14.45 | 271.976 |
| Mia_100 | 3010.0 | 5.614 | 12.299 | 0.825 | 2.079 | 2.587 | 3.556 | 211.245 |
| Mia_200 | 3232.0 | 10.731 | 23.187 | 0.75 | 2.875 | 3.987 | 7.175 | 241.359 |
| Mia_300 | 3232.0 | 18.731 | 39.479 | 1.033 | 3.491 | 5.461 | 12.515 | 274.812 |
| Mia_wash | 4732.0 | 5.017 | 11.733 | 0.478 | 2.42 | 2.958 | 3.98 | 211.304 |
| P10_100 | 8226.0 | 57.826 | 18.417 | 2.416 | 44.851 | 56.396 | 68.098 | 183.769 |
| P10_200 | 8226.0 | 116.058 | 41.896 | 3.631 | 97.607 | 120.635 | 144.736 | 262.384 |
| P10_300 | 6857.0 | 148.842 | 50.68 | 2.29 | 110.504 | 137.728 | 177.635 | 299.611 |
| P10_wash | 5935.0 | 172.491 | 40.544 | 3.954 | 144.878 | 162.806 | 203.494 | 294.754 |

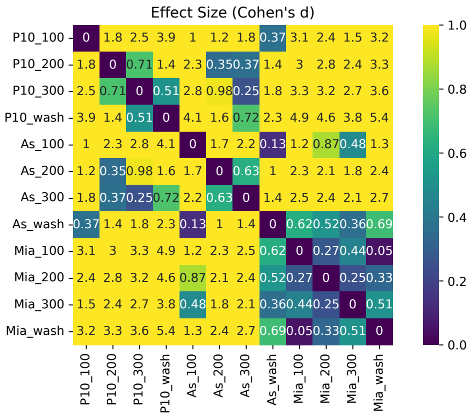


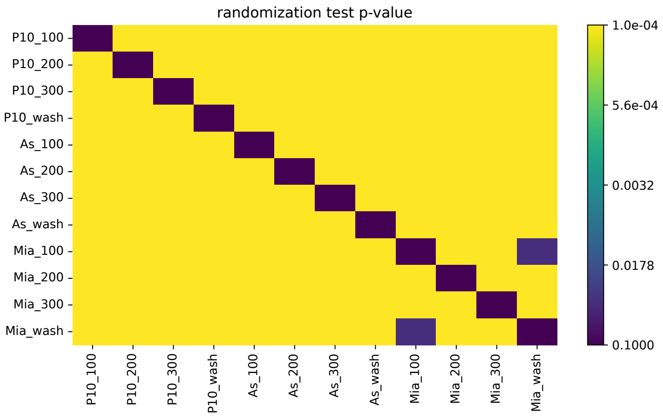

#### FigS3B_MinSpeed

| Condition | count | mean | std | min | 25% | 50% | 75% | max |
|---|---|---|---|---|---|---|---|---|
| As_100 | 7329.0 | 5.497 | 9.709 | 0.0 | 0.0 | 0.661 | 6.058 | 72.426 |
| As_200 | 7043.0 | 31.388 | 32.12 | 0.0 | 3.134 | 19.294 | 55.05 | 157.902 |
| As_300 | 6591.0 | 45.23 | 48.054 | 0.0 | 6.441 | 26.209 | 73.428 | 241.697 |
| As_wash | 5325.0 | 12.946 | 30.718 | 0.0 | 0.0 | 0.0 | 0.417 | 194.491 |
| Mia_100 | 3010.0 | 0.497 | 4.068 | 0.0 | 0.0 | 0.0 | 0.0 | 101.851 |
| Mia_200 | 3232.0 | 1.451 | 8.88 | 0.0 | 0.0 | 0.0 | 0.0 | 155.843 |
| Mia_300 | 2416.0 | 5.22 | 21.009 | 0.0 | 0.0 | 0.0 | 0.606 | 182.49 |
| Mia_wash | 4732.0 | 0.315 | 4.095 | 0.0 | 0.0 | 0.0 | 0.0 | 116.312 |
| P10_100 | 8226.0 | 14.317 | 10.139 | 0.0 | 6.587 | 12.012 | 20.561 | 97.339 |
| P10_200 | 8213.0 | 45.944 | 27.976 | 0.0 | 24.919 | 44.9 | 64.532 | 179.647 |
| P10_300 | 6838.0 | 68.542 | 43.069 | 0.0 | 37.988 | 56.854 | 90.074 | 231.871 |
| P10_wash | 5935.0 | 83.349 | 34.35 | 0.0 | 61.463 | 81.379 | 103.76 | 209.856 |

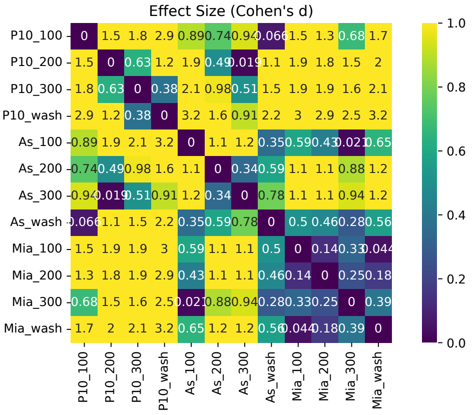


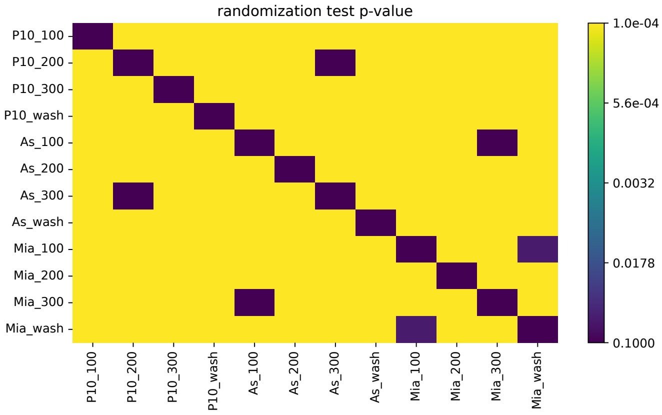


#### FigS3D_MaxSpeed

| Condition | count | mean | std | min | 25% | 50% | 75% | max |
|---|---|---|---|---|---|---|---|---|
| IL_Mono_100 | 7031.0 | 85.504 | 70.49 | 4.557 | 34.965 | 68.359 | 116.394 | 372.275 |
| IL_Mono_200 | 7031.0 | 128.151 | 91.008 | 4.686 | 49.615 | 118.663 | 179.527 | 374.807 |
| IL_Mono_300 | 5565.0 | 157.266 | 93.257 | 5.126 | 95.229 | 155.744 | 221.221 | 374.89 |
| IL_Mono_wash | 8188.0 | 186.48 | 91.621 | 5.599 | 142.064 | 194.591 | 239.427 | 374.905 |
| IL_Neut_100 | 4451.0 | 58.371 | 64.929 | 4.503 | 15.542 | 28.972 | 79.14 | 372.618 |
| IL_Neut_200 | 4451.0 | 87.558 | 90.888 | 3.574 | 17.314 | 40.137 | 146.026 | 374.738 |
| IL_Neut_300 | 3044.0 | 119.623 | 100.299 | 3.574 | 22.49 | 110.788 | 193.663 | 374.632 |
| IL_Neut_wash | 4976.0 | 92.395 | 95.881 | 3.823 | 18.035 | 37.421 | 163.646 | 374.969 |
| Mono_100 | 10947.0 | 96.95 | 70.397 | 5.344 | 55.819 | 75.19 | 107.74 | 374.822 |
| Mono_200 | 10947.0 | 153.287 | 84.359 | 5.794 | 97.58 | 139.664 | 193.258 | 374.685 |
| Mono_300 | 8045.0 | 200.241 | 89.878 | 5.692 | 139.36 | 199.245 | 263.687 | 374.737 |
| Mono_wash | 12548.0 | 211.93 | 72.561 | 6.706 | 161.808 | 206.574 | 257.311 | 375.0 |
| Neut_100 | 10063.0 | 70.893 | 65.099 | 3.531 | 19.288 | 59.657 | 94.841 | 374.582 |
| Neut_200 | 10063.0 | 112.52 | 92.403 | 4.297 | 22.929 | 98.434 | 175.532 | 373.919 |
| Neut_300 | 7028.0 | 157.315 | 99.293 | 4.49 | 48.301 | 171.652 | 230.086 | 374.507 |
| Neut_wash | 12904.0 | 151.909 | 104.004 | 3.727 | 33.535 | 168.838 | 230.783 | 374.971 |

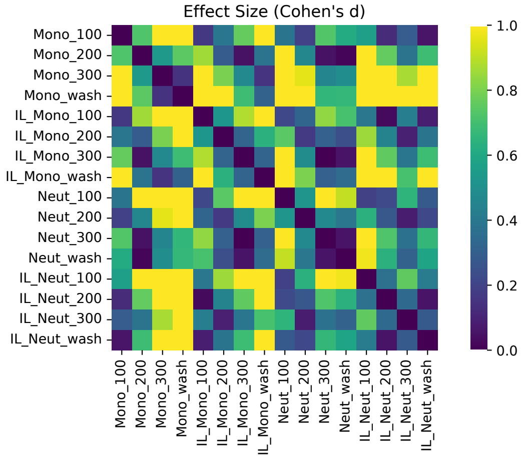


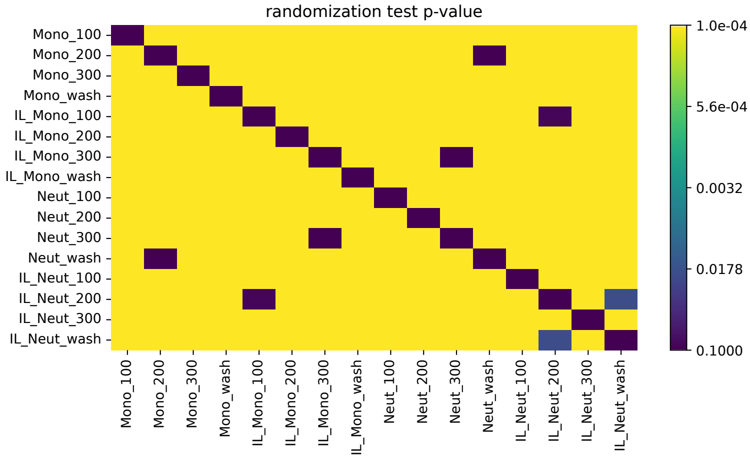

### FigS3D_MeanSpeed

| Condition | count | mean | std | min | 25% | 50% | 75% | max |
|---|---|---|---|---|---|---|---|---|
| IL_Mono_100 | 7031.0 | 22.915 | 15.288 | 0.987 | 4.691 | 24.659 | 32.959 | 105.231 |
| IL_Mono_200 | 7031.0 | 44.755 | 33.825 | 1.398 | 9.897 | 46.076 | 71.538 | 152.487 |
| IL_Mono_300 | 5565.0 | 61.084 | 46.133 | 1.779 | 11.084 | 58.484 | 91.303 | 241.002 |
| IL_Mono_wash | 8188.0 | 64.467 | 50.319 | 1.24 | 12.235 | 64.27 | 108.968 | 217.073 |
| IL_Neut_100 | 4451.0 | 9.304 | 11.657 | 0.938 | 2.683 | 3.664 | 8.905 | 79.305 |
| IL_Neut_200 | 4451.0 | 14.63 | 23.069 | 0.841 | 2.942 | 4.138 | 11.148 | 135.763 |
| IL_Neut_300 | 3044.0 | 24.839 | 39.331 | 0.841 | 3.445 | 5.348 | 22.883 | 249.186 |
| IL_Neut_wash | 4976.0 | 12.194 | 23.81 | 1.236 | 3.244 | 4.129 | 6.786 | 197.982 |
| Mono_100 | 10947.0 | 32.752 | 14.852 | 1.509 | 24.65 | 31.434 | 39.549 | 133.349 |
| Mono_200 | 10947.0 | 61.623 | 31.833 | 1.574 | 32.475 | 62.927 | 82.732 | 181.858 |
| Mono_300 | 8045.0 | 85.635 | 45.198 | 1.438 | 56.769 | 77.701 | 117.985 | 260.422 |
| Mono_wash | 12548.0 | 108.633 | 40.302 | 2.335 | 85.939 | 108.369 | 135.152 | 259.621 |
| Neut_100 | 10063.0 | 17.168 | 17.301 | 1.029 | 3.19 | 6.242 | 30.796 | 95.352 |
| Neut_200 | 10063.0 | 32.46 | 36.43 | 0.72 | 3.554 | 9.478 | 61.961 | 183.925 |
| Neut_300 | 7028.0 | 56.367 | 55.82 | 0.72 | 4.774 | 38.719 | 101.08 | 246.327 |
| Neut_wash | 12904.0 | 43.192 | 49.692 | 1.077 | 4.416 | 10.437 | 86.804 | 243.824 |

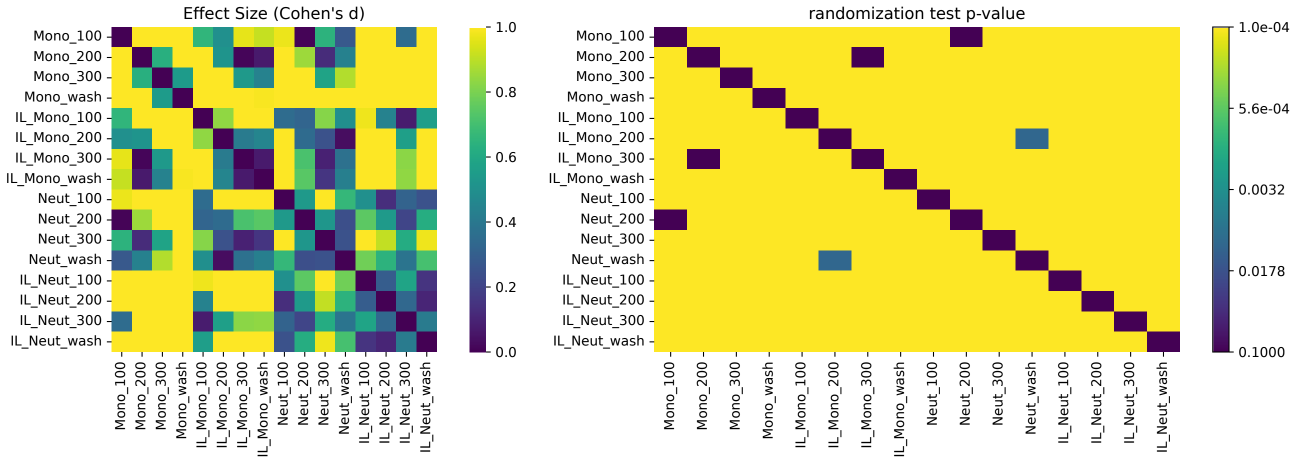

### FigS3D_MinSpeed

| Condition | count | mean | std | min | 25% | 50% | 75% | max |
|---|---|---|---|---|---|---|---|---|
| IL_Mono_100 | 7031.0 | 2.065 | 3.619 | 0.0 | 0.0 | 0.65 | 2.588 | 34.076 |
| IL_Mono_200 | 6655.0 | 11.854 | 14.069 | 0.0 | 0.0 | 5.495 | 20.303 | 90.372 |
| IL_Mono_300 | 4529.0 | 23.207 | 23.505 | 0.0 | 2.162 | 17.907 | 34.104 | 160.152 |
| IL_Mono_wash | 8188.0 | 21.022 | 28.951 | 0.0 | 0.0 | 0.317 | 42.966 | 151.713 |
| IL_Neut_100 | 4451.0 | 0.387 | 1.692 | 0.0 | 0.0 | 0.0 | 0.0 | 32.235 |
| IL_Neut_200 | 3232.0 | 2.304 | 6.597 | 0.0 | 0.0 | 0.0 | 0.0 | 65.492 |
| IL_Neut_300 | 1522.0 | 9.836 | 20.44 | 0.0 | 0.0 | 0.0 | 9.364 | 156.179 |
| IL_Neut_wash | 4976.0 | 1.557 | 7.76 | 0.0 | 0.0 | 0.0 | 0.0 | 107.98 |
| Mono_100 | 10947.0 | 6.814 | 6.347 | 0.0 | 2.013 | 5.344 | 10.032 | 52.524 |
| Mono_200 | 10537.0 | 21.475 | 18.287 | 0.0 | 6.369 | 17.42 | 32.298 | 114.009 |
| Mono_300 | 7679.0 | 33.456 | 27.987 | 0.0 | 13.479 | 24.539 | 48.114 | 164.056 |
| Mono_wash | 12548.0 | 50.246 | 30.161 | 0.0 | 32.696 | 48.404 | 69.389 | 186.324 |
| Neut_100 | 10063.0 | 2.223 | 5.252 | 0.0 | 0.0 | 0.0 | 1.387 | 50.766 |
| Neut_200 | 7844.0 | 7.966 | 14.893 | 0.0 | 0.0 | 0.333 | 8.373 | 117.752 |
| Neut_300 | 4858.0 | 23.077 | 30.553 | 0.0 | 0.0 | 5.833 | 38.843 | 181.094 |
| Neut_wash | 12904.0 | 9.859 | 20.487 | 0.0 | 0.0 | 0.0 | 7.519 | 157.609 |

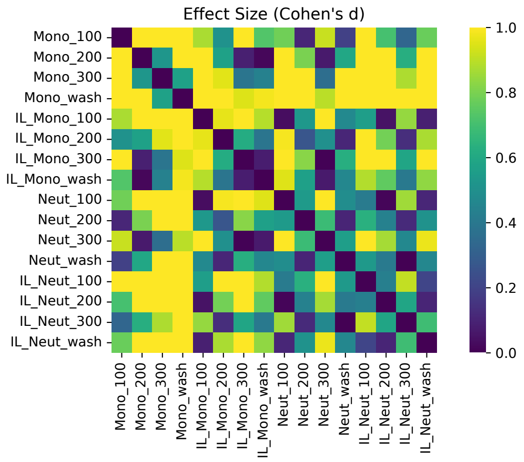


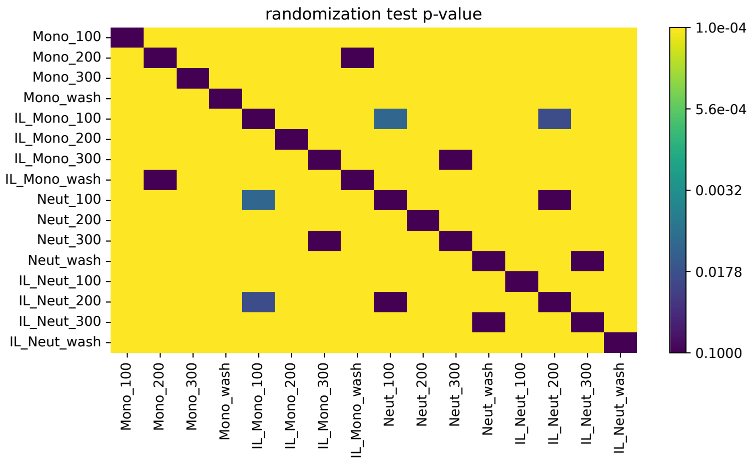

FigS5A

| Condition | count | mean | std | min | 25% | 50% | 75% | max |
|---|---|---|---|---|---|---|---|---|
| End_As_IL1b_junc | 84.0 | 5.873 | 3.395 | 0.65 | 3.248 | 5.699 | 8.216 | 14.699 |
| End_As_IL1b_nuc | 84.0 | 9.088 | 7.345 | 0.0 | 1.837 | 9.23 | 13.599 | 25.985 |
| End_As_ctrl_junc | 334.0 | 5.621 | 3.117 | 0.0 | 3.788 | 5.297 | 7.263 | 21.16 |
| End_As_ctrl_nuc | 334.0 | 10.023 | 7.645 | 0.0 | 2.342 | 9.787 | 16.201 | 28.583 |
| End_Mia_IL1b_junc | 191.0 | 5.217 | 3.884 | 0.0 | 2.598 | 4.684 | 7.146 | 29.634 |
| End_Mia_IL1b_nuc | 191.0 | 9.756 | 8.793 | 0.0 | 1.376 | 9.187 | 13.734 | 40.931 |
| End_Mia_ctrl_junc | 395.0 | 5.703 | 5.612 | 0.0 | 2.342 | 4.547 | 7.604 | 54.849 |
| End_Mia_ctrl_nuc | 395.0 | 9.851 | 8.472 | 0.0 | 2.752 | 9.301 | 13.925 | 68.321 |
| End_Mono_IL1b_junc | 1105.0 | 4.996 | 3.581 | 0.0 | 2.342 | 4.358 | 6.905 | 21.526 |
| End_Mono_IL1b_nuc | 1105.0 | 7.083 | 6.135 | 0.0 | 1.949 | 5.989 | 11.043 | 32.364 |
| End_Mono_ctrl_junc | 359.0 | 5.222 | 3.884 | 0.0 | 2.598 | 4.358 | 6.997 | 23.709 |
| End_Mono_ctrl_nuc | 359.0 | 8.17 | 6.723 | 0.0 | 3.373 | 6.625 | 11.519 | 34.582 |
| End_Neu_IL1b_junc | 1139.0 | 5.731 | 4.429 | 0.0 | 2.598 | 4.593 | 8.14 | 33.761 |
| End_Neu_IL1b_nuc | 1139.0 | 8.128 | 7.151 | 0.0 | 2.678 | 6.496 | 12.17 | 39.966 |
| End_Neu_ctrl_junc | 1710.0 | 4.953 | 3.996 | 0.0 | 1.949 | 4.109 | 6.997 | 23.422 |
| End_Neu_ctrl_nuc | 1710.0 | 8.404 | 6.947 | 0.0 | 3.248 | 7.175 | 12.205 | 45.366 |
| landing_As_IL1b_junct | 84.0 | 5.061 | 3.305 | 0.0 | 2.342 | 4.547 | 6.928 | 13.657 |
| landing_As_IL1b_nuc | 84.0 | 8.631 | 7.124 | 0.0 | 2.598 | 7.576 | 12.996 | 30.135 |
| landing_As_ctrl_junct | 334.0 | 5.212 | 3.463 | 0.0 | 2.678 | 4.947 | 6.997 | 21.437 |
| landing_As_ctrl_nuc | 334.0 | 10.365 | 7.177 | 0.0 | 4.682 | 10.535 | 15.441 | 27.933 |
| landing_Mia_IL1b_junct | 191.0 | 5.485 | 4.348 | 0.0 | 2.002 | 4.547 | 8.382 | 28.59 |
| landing_Mia_IL1b_nuc | 191.0 | 8.959 | 7.935 | 0.0 | 1.893 | 8.445 | 12.629 | 40.553 |
| landing_Mia_ctrl_junct | 395.0 | 5.29 | 4.905 | 0.0 | 2.054 | 4.358 | 7.491 | 56.262 |
| landing_Mia_ctrl_nuc | 395.0 | 9.75 | 8.5 | 0.0 | 4.109 | 8.74 | 13.384 | 69.192 |
| landing_Mono_IL1b_junct | 1105.0 | 5.068 | 3.697 | 0.0 | 2.342 | 4.547 | 7.263 | 20.949 |
| landing_Mono_IL1b_nuc | 1105.0 | 7.158 | 6.084 | 0.0 | 1.949 | 5.989 | 11.043 | 32.772 |
| landing_Mono_ctrl_junct | 359.0 | 5.237 | 3.877 | 0.0 | 2.598 | 4.593 | 6.951 | 23.709 |
| landing_Mono_ctrl_nuc | 359.0 | 7.914 | 6.268 | 0.0 | 3.587 | 7.027 | 11.053 | 32.875 |
| landing_Neu_IL1b_junct | 1139.0 | 5.772 | 4.556 | 0.0 | 2.054 | 4.684 | 8.507 | 28.494 |
| landing_Neu_IL1b_nuc | 1139.0 | 8.159 | 7.375 | 0.0 | 1.949 | 6.496 | 12.445 | 45.236 |
| landing_Neu_ctrl_junct | 1710.0 | 4.889 | 3.947 | 0.0 | 1.949 | 3.898 | 6.997 | 29.748 |
| landing_Neu_ctrl_nuc | 1710.0 | 8.336 | 6.882 | 0.0 | 2.756 | 7.263 | 12.326 | 40.683 |

FigS5B

| Condition | count | mean | std | min | 25% | 50% | 75% | max |
|---|---|---|---|---|---|---|---|---|
| arrest_As_IL1b_junct | 84.0 | 5.442 | 3.223 | 0.0 | 3.248 | 5.01 | 7.299 | 13.704 |
| arrest_As_IL1b_nuc | 84.0 | 8.667 | 7.443 | 0.0 | 2.028 | 7.361 | 12.562 | 26.634 |
| arrest_As_ctrl_junct | 334.0 | 5.596 | 3.419 | 0.0 | 3.498 | 5.237 | 7.263 | 23.041 |
| arrest_As_ctrl_nuc | 334.0 | 9.895 | 7.367 | 0.0 | 3.264 | 9.744 | 15.383 | 27.933 |
| arrest_Mia_IL1b_junct | 191.0 | 5.354 | 4.113 | 0.0 | 2.342 | 4.594 | 7.836 | 28.59 |
| arrest_Mia_IL1b_nuc | 191.0 | 9.526 | 8.551 | 0.0 | 1.376 | 8.836 | 13.353 | 40.553 |
| arrest_Mia_ctrl_junct | 395.0 | 5.3 | 5.006 | 0.0 | 2.002 | 4.594 | 7.146 | 56.699 |
| arrest_Mia_ctrl_nuc | 395.0 | 9.614 | 8.75 | 0.0 | 2.598 | 8.667 | 13.773 | 69.107 |
| arrest_Mono_IL1b_junct | 1105.0 | 5.068 | 3.644 | 0.0 | 2.342 | 4.358 | 7.146 | 21.526 |
| arrest_Mono_IL1b_nuc | 1105.0 | 7.082 | 6.132 | 0.0 | 1.949 | 5.989 | 11.043 | 32.772 |
| arrest_Mono_ctrl_junct | 359.0 | 5.251 | 3.914 | 0.0 | 2.47 | 4.547 | 6.997 | 23.824 |
| arrest_Mono_ctrl_nuc | 359.0 | 8.089 | 6.608 | 0.0 | 3.248 | 6.688 | 11.428 | 33.156 |
| arrest_Neu_IL1b_junct | 1139.0 | 5.746 | 4.517 | 0.0 | 2.054 | 4.947 | 8.457 | 33.003 |
| arrest_Neu_IL1b_nuc | 1139.0 | 8.155 | 7.274 | 0.0 | 2.054 | 6.688 | 12.496 | 39.069 |
| arrest_Neu_ctrl_junct | 1710.0 | 4.978 | 4.061 | 0.0 | 1.949 | 4.108 | 6.997 | 29.748 |
| arrest_Neu_ctrl_nuc | 1710.0 | 8.474 | 6.907 | 0.0 | 2.905 | 7.604 | 12.725 | 42.894 |
| landing_As_IL1b_junct | 84.0 | 5.061 | 3.305 | 0.0 | 2.342 | 4.547 | 6.928 | 13.657 |
| landing_As_IL1b_nuc | 84.0 | 8.631 | 7.124 | 0.0 | 2.598 | 7.576 | 12.996 | 30.135 |
| landing_As_ctrl_junct | 334.0 | 5.212 | 3.463 | 0.0 | 2.678 | 4.947 | 6.997 | 21.437 |
| landing_As_ctrl_nuc | 334.0 | 10.365 | 7.177 | 0.0 | 4.682 | 10.535 | 15.441 | 27.933 |
| landing_Mia_IL1b_junct | 191.0 | 5.485 | 4.348 | 0.0 | 2.002 | 4.547 | 8.382 | 28.59 |
| landing_Mia_IL1b_nuc | 191.0 | 8.959 | 7.935 | 0.0 | 1.893 | 8.445 | 12.629 | 40.553 |
| landing_Mia_ctrl_junct | 395.0 | 5.29 | 4.905 | 0.0 | 2.054 | 4.358 | 7.491 | 56.262 |
| landing_Mia_ctrl_nuc | 395.0 | 9.75 | 8.5 | 0.0 | 4.109 | 8.74 | 13.384 | 69.192 |
| landing_Mono_IL1b_junct | 1105.0 | 5.068 | 3.697 | 0.0 | 2.342 | 4.547 | 7.263 | 20.949 |
| landing_Mono_IL1b_nuc | 1105.0 | 7.158 | 6.084 | 0.0 | 1.949 | 5.989 | 11.043 | 32.772 |
| landing_Mono_ctrl_junct | 359.0 | 5.237 | 3.877 | 0.0 | 2.598 | 4.593 | 6.951 | 23.709 |
| landing_Mono_ctrl_nuc | 359.0 | 7.914 | 6.268 | 0.0 | 3.587 | 7.027 | 11.053 | 32.875 |
| landing_Neu_IL1b_junct | 1139.0 | 5.772 | 4.556 | 0.0 | 2.054 | 4.684 | 8.507 | 28.494 |
| landing_Neu_IL1b_nuc | 1139.0 | 8.159 | 7.375 | 0.0 | 1.949 | 6.496 | 12.445 | 45.236 |
| landing_Neu_ctrl_junct | 1710.0 | 4.889 | 3.947 | 0.0 | 1.949 | 3.898 | 6.997 | 29.748 |
| landing_Neu_ctrl_nuc | 1710.0 | 8.336 | 6.882 | 0.0 | 2.756 | 7.263 | 12.326 | 40.683 |

### FigS6B

| Condition | count | mean | std | min | 25% | 50% | 75% | max |
|---|---|---|---|---|---|---|---|---|
| ASPC1 | 40.0 | 53.623 | 43.884 | 4.658 | 26.897 | 42.002 | 73.966 | 188.118 |
| M-C simu | 192.0 | -2.997 | 6.944 | -50.0 | -1.921 | -1.726 | -1.642 | -1.168 |
| Monocytes | 40.0 | 31.292 | 26.843 | 1.67 | 13.918 | 25.034 | 39.004 | 129.097 |
| Neutrophils | 48.0 | 13.243 | 8.685 | 0.561 | 6.267 | 13.025 | 17.544 | 44.164 |
| miapaca | 32.0 | 23.324 | 14.368 | 5.849 | 11.523 | 21.024 | 33.058 | 62.035 |

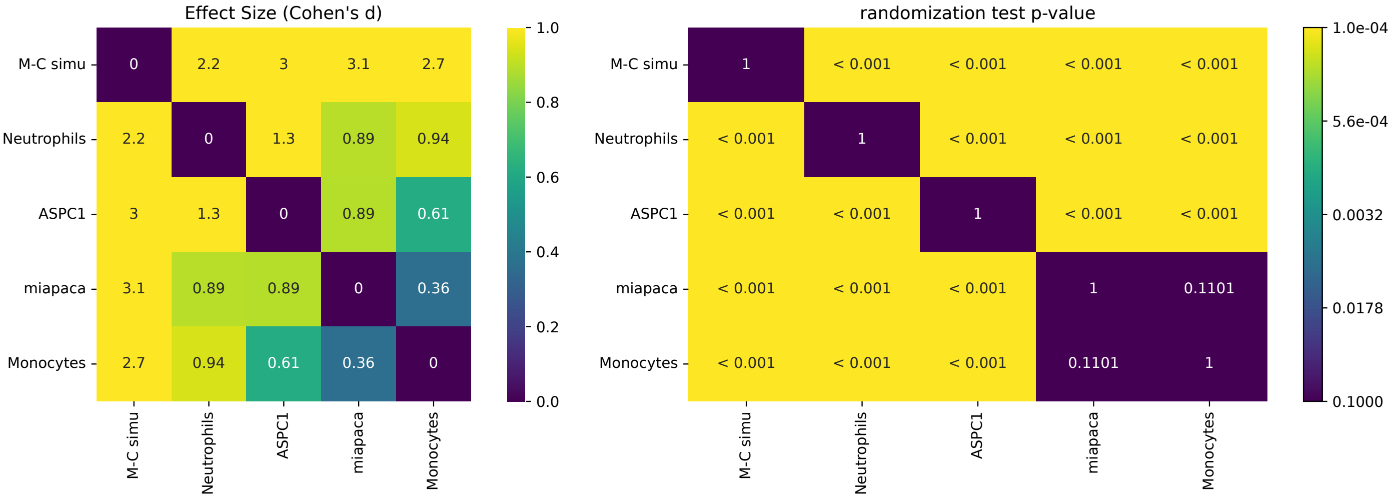


### FigS7A

| Condition | count | mean | std | min | 25% | 50% | 75% | max |
|---|---|---|---|---|---|---|---|---|
| As | 3.0 | 0.086 | 0.02 | 0.069 | 0.075 | 0.082 | 0.095 | 0.108 |
| Mia | 3.0 | 1.0 | 0.193 | 0.777 | 0.941 | 1.106 | 1.112 | 1.117 |
| P10 | 3.0 | 0.015 | 0.013 | 0.0 | 0.009 | 0.018 | 0.022 | 0.026 |

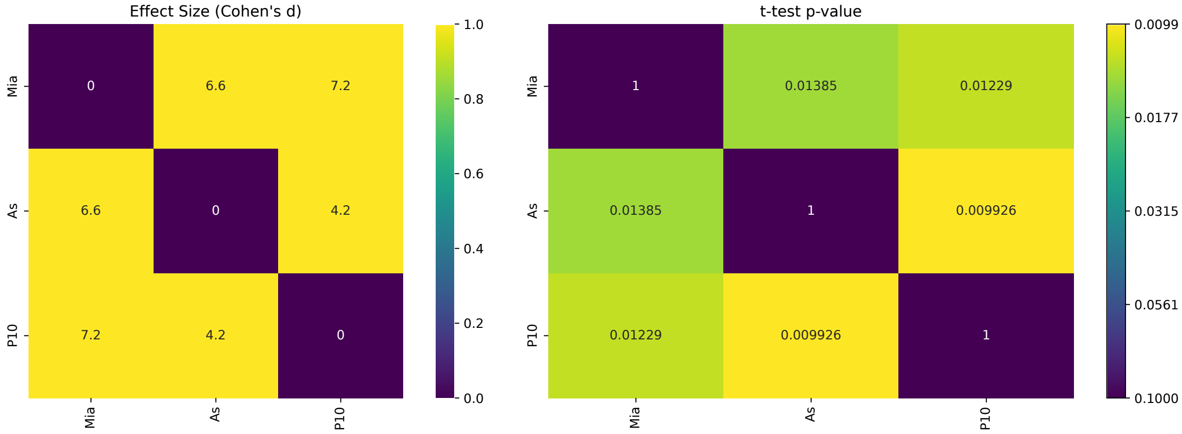

#### FigS7B

| Condition | count | mean | std | min | 25% | 50% | 75% | max |
|---|---|---|---|---|---|---|---|---|
| siCD44 #1 | 8.0 | 0.489 | 0.102 | 0.36 | 0.425 | 0.47 | 0.562 | 0.66 |
| siCD44 #2 | 2.0 | 0.485 | 0.035 | 0.46 | 0.473 | 0.485 | 0.498 | 0.51 |
| siCD44 #3 | 8.0 | 0.489 | 0.234 | 0.19 | 0.33 | 0.475 | 0.548 | 0.96 |
| siCtrl | 8.0 | 1.0 | 0.213 | 0.73 | 0.915 | 0.97 | 1.0 | 1.47 |

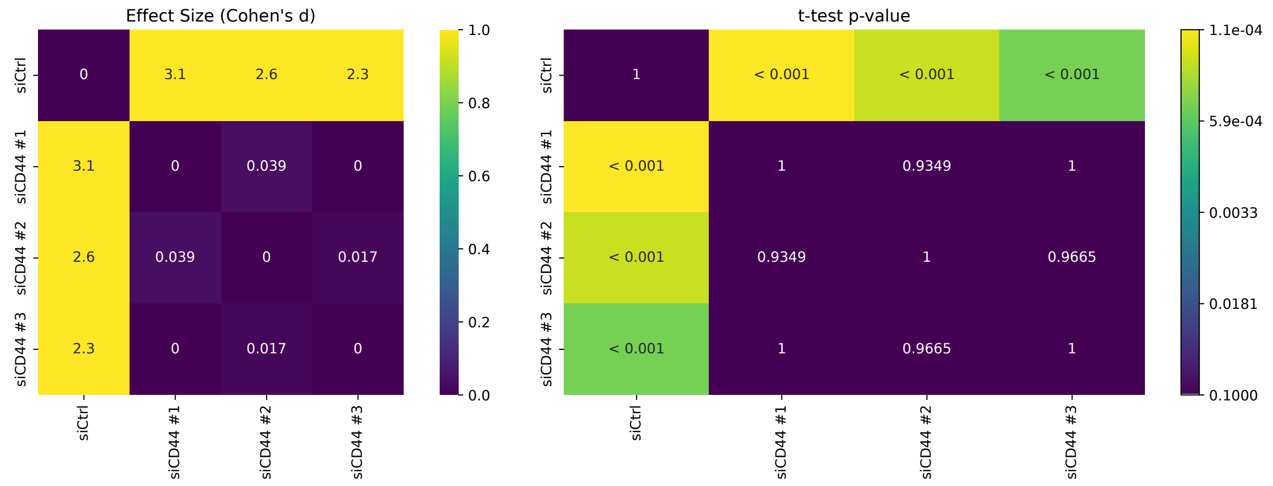


#### FigS7C

| Condition | count | mean | std | min | 25% | 50% | 75% | max |
|---|---|---|---|---|---|---|---|---|
| siCD44 #1 | 3.0 | 0.538 | 0.368 | 0.131 | 0.384 | 0.636 | 0.741 | 0.846 |
| siCD44 #3 | 3.0 | 0.431 | 0.256 | 0.153 | 0.318 | 0.482 | 0.57 | 0.657 |
| siCtrl | 3.0 | 1.0 | 0.0 | 1.0 | 1.0 | 1.0 | 1.0 | 1.0 |

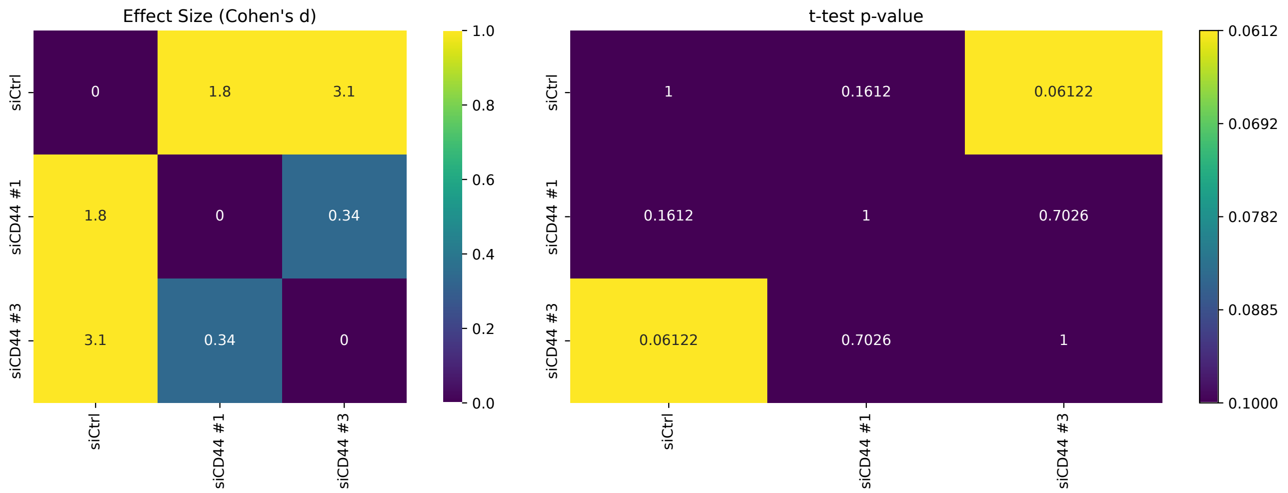

### FigS7D

| Condition | count | mean | std | min | 25% | 50% | 75% | max |
|---|---|---|---|---|---|---|---|---|
| siCD44 #1 | 55.0 | 0.917 | 0.136 | 0.458 | 0.871 | 0.929 | 1.011 | 1.096 |
| siCD44 #3 | 60.0 | 0.883 | 0.132 | 0.438 | 0.862 | 0.898 | 0.964 | 1.061 |
| siCtrl | 53.0 | 1.0 | 0.195 | 0.516 | 0.813 | 1.043 | 1.16 | 1.357 |

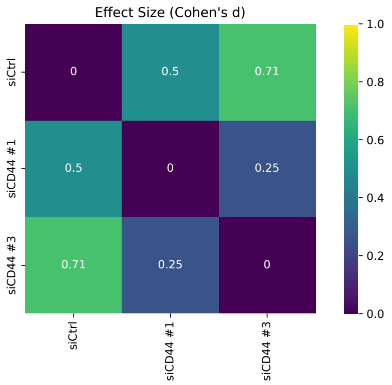


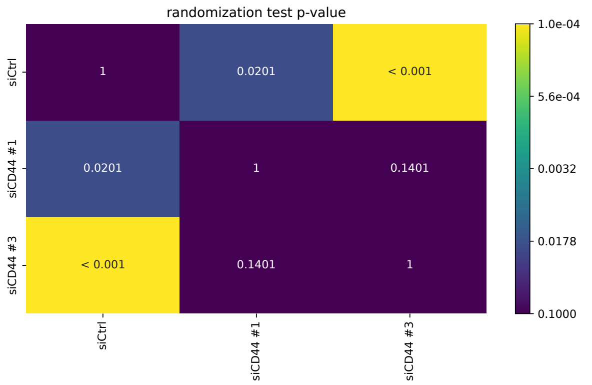


### FigS7G

| Condition | count | mean | std | min | 25% | 50% | 75% | max |
|---|---|---|---|---|---|---|---|---|
| Mia si2 200 | 4.0 | 0.299 | 0.304 | 0.041 | 0.043 | 0.262 | 0.519 | 0.632 |
| as ctrl 200 | 2.0 | 0.32 | 0.319 | 0.094 | 0.207 | 0.32 | 0.432 | 0.545 |
| as s1 200 | 2.0 | 0.548 | 0.363 | 0.292 | 0.42 | 0.548 | 0.676 | 0.805 |
| as si2 200 | 2.0 | 0.525 | 0.145 | 0.422 | 0.474 | 0.525 | 0.576 | 0.628 |
| as si3 200 | 2.0 | 0.323 | 0.179 | 0.197 | 0.26 | 0.323 | 0.387 | 0.45 |
| mia ctrl 200 | 4.0 | 0.495 | 0.076 | 0.429 | 0.431 | 0.49 | 0.554 | 0.572 |
| mia s1 200 | 4.0 | 0.259 | 0.169 | 0.13 | 0.135 | 0.208 | 0.332 | 0.492 |
| mia si3 200 | 4.0 | 0.444 | 0.136 | 0.247 | 0.426 | 0.487 | 0.505 | 0.556 |

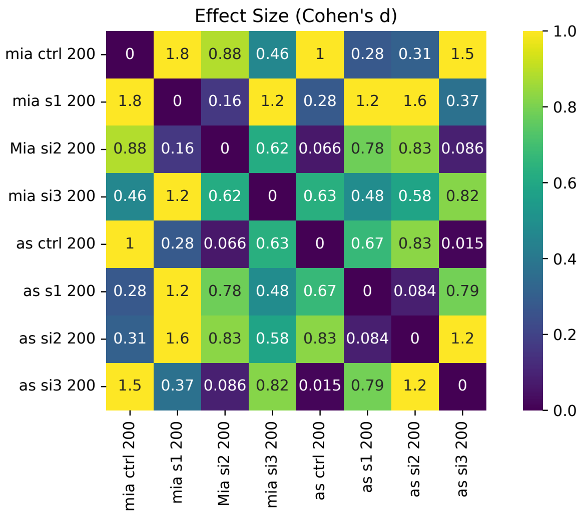


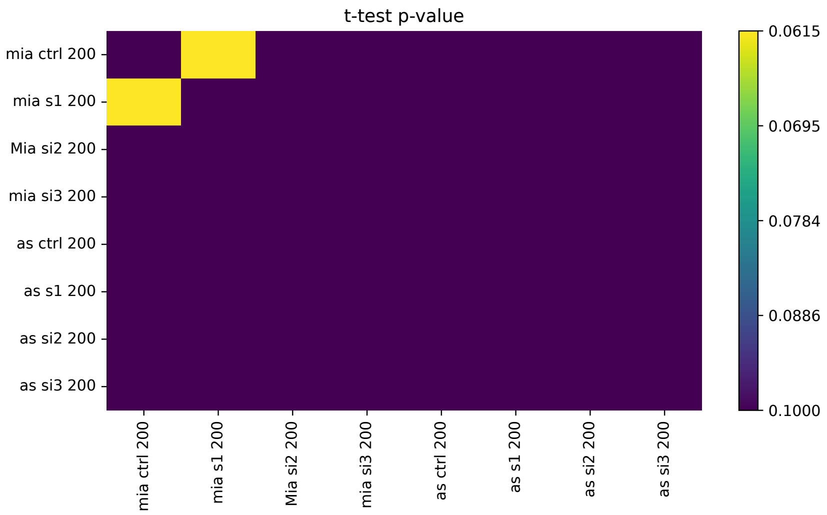

#### FigS7H

| Condition | count | mean | std | min | 25% | 50% | 75% | max |
|---|---|---|---|---|---|---|---|---|
| As si2 | 8.0 | 72.337 | 34.255 | 33.111 | 37.238 | 77.887 | 96.407 | 117.764 |
| as ctrl | 8.0 | 67.922 | 15.662 | 41.423 | 58.26 | 69.021 | 76.803 | 92.511 |
| as si1 | 8.0 | 65.383 | 35.767 | 23.948 | 33.316 | 65.552 | 87.167 | 119.709 |
| as si3 | 8.0 | 68.197 | 26.911 | 32.204 | 47.719 | 70.084 | 81.909 | 111.763 |
| mia ctrl | 16.0 | 50.859 | 18.421 | 27.791 | 37.981 | 49.055 | 60.388 | 93.102 |
| mia si1 | 16.0 | 66.115 | 28.46 | 27.043 | 44.295 | 61.368 | 84.851 | 127.779 |
| mia si2 | 16.0 | 71.738 | 28.063 | 37.107 | 48.291 | 69.34 | 82.651 | 142.838 |
| mia si3 | 16.0 | 52.201 | 19.12 | 21.894 | 40.778 | 48.332 | 65.34 | 87.889 |

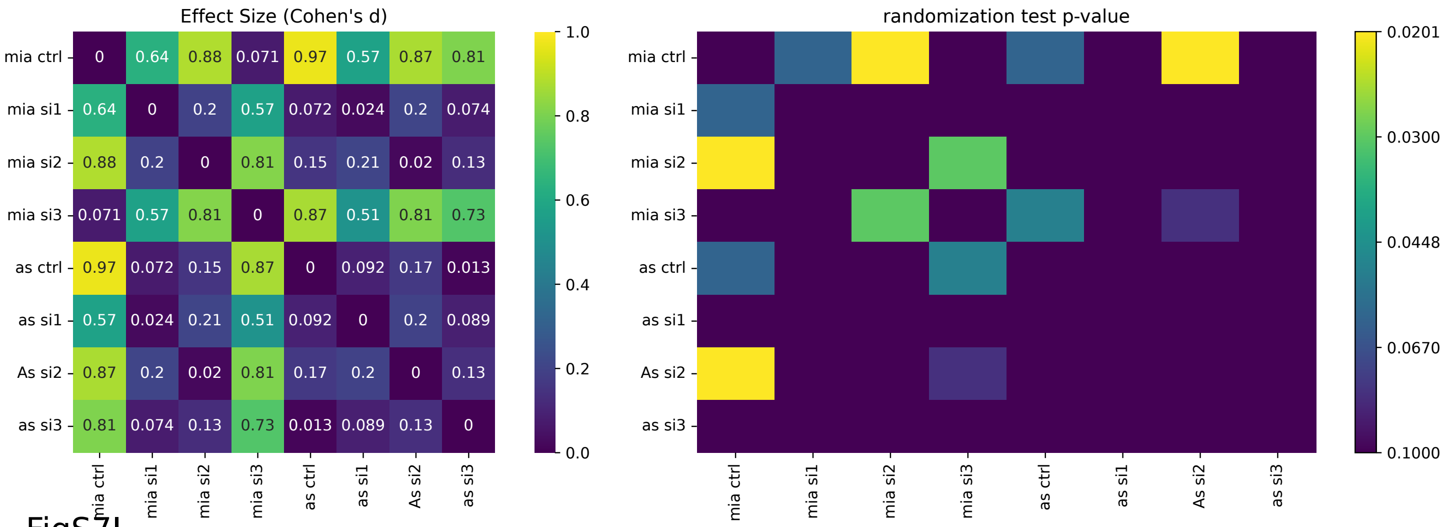


#### FigS7I

| Condition | count | mean | std | min | 25% | 50% | 75% | max |
|---|---|---|---|---|---|---|---|---|
| As si2 | 458.0 | 184.524 | 175.019 | 0.22 | 56.266 | 142.477 | 250.769 | 1284.674 |
| as ctrl | 374.0 | 174.055 | 166.984 | 1.499 | 55.88 | 138.016 | 242.072 | 1395.54 |
| as si1 | 474.0 | 167.918 | 176.434 | 0.0 | 47.477 | 123.145 | 236.722 | 1456.752 |
| as si3 | 331.0 | 159.037 | 158.152 | 0.0 | 36.588 | 107.604 | 226.863 | 869.059 |
| mia ctrl | 1248.0 | 170.236 | 177.693 | 0.0 | 46.399 | 115.358 | 246.921 | 1428.748 |
| mia si1 | 793.0 | 144.883 | 148.619 | 0.0 | 38.133 | 99.501 | 212.603 | 1090.475 |
| mia si2 | 632.0 | 153.663 | 158.198 | 0.0 | 40.725 | 109.841 | 211.414 | 1195.262 |
| mia si3 | 859.0 | 148.418 | 164.212 | 0.0 | 30.737 | 90.128 | 214.732 | 1128.014 |

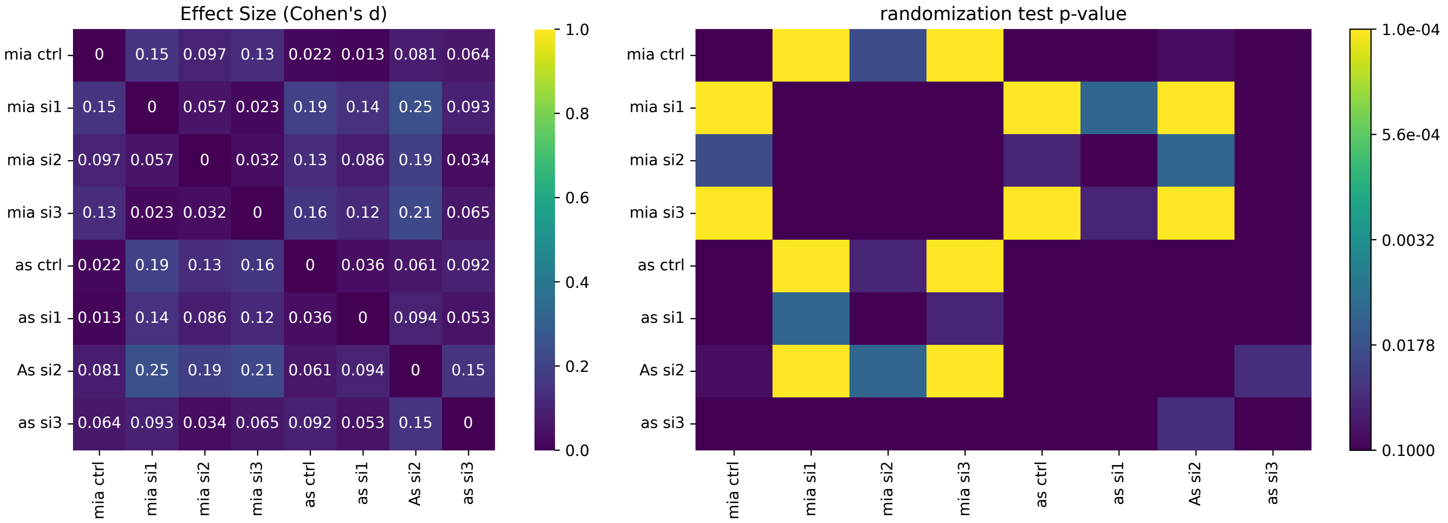

### FigS7J

| Condition | count | mean | std | min | 25% | 50% | 75% | max |
|---|---|---|---|---|---|---|---|---|
| As si1 | 474.0 | 34.477 | 29.794 | 0.0 | 10.0 | 26.5 | 50.75 | 151.0 |
| As si2 | 458.0 | 39.924 | 32.144 | 0.0 | 13.0 | 33.5 | 59.0 | 166.0 |
| As si3 | 331.0 | 35.776 | 33.493 | 0.0 | 9.0 | 27.0 | 53.0 | 173.0 |
| as sctrl | 374.0 | 38.147 | 30.9 | 0.0 | 12.25 | 31.0 | 55.0 | 159.0 |
| mai ctrl | 1248.0 | 36.34 | 31.972 | 0.0 | 11.0 | 26.0 | 56.0 | 168.0 |
| mia s2 | 632.0 | 33.707 | 31.118 | 0.0 | 9.0 | 24.5 | 49.25 | 176.0 |
| mia si1 | 793.0 | 32.182 | 29.857 | 0.0 | 9.0 | 23.0 | 49.0 | 180.0 |
| mia si3 | 859.0 | 33.146 | 32.5 | 0.0 | 8.0 | 20.0 | 49.0 | 156.0 |

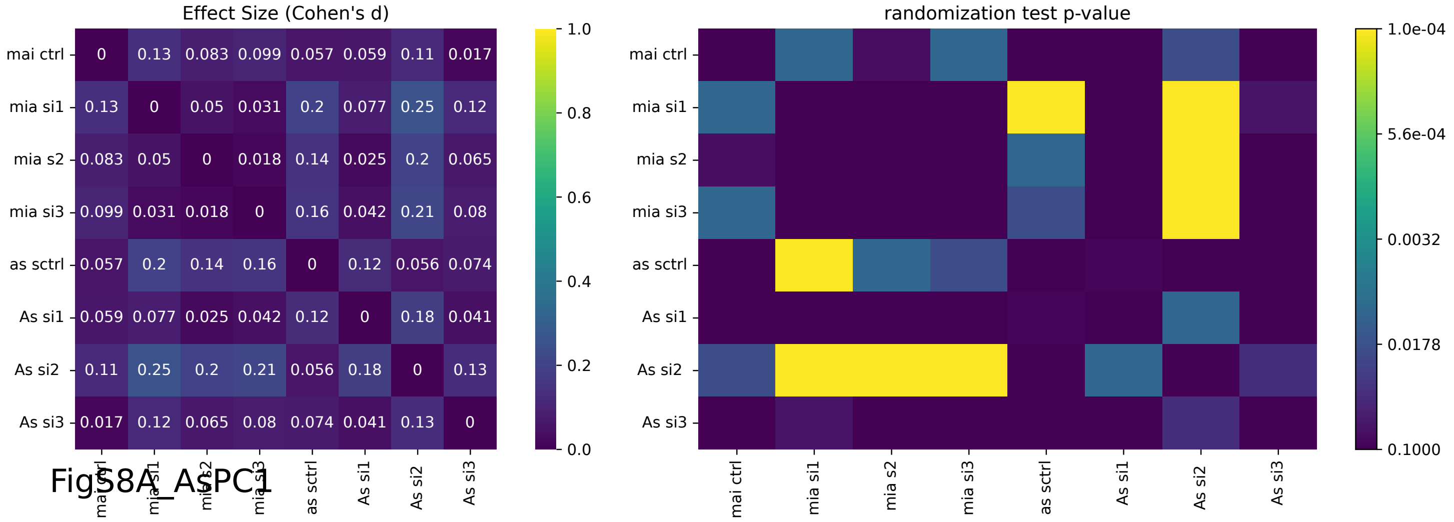


### FigS8A_ASPC1

| Condition | count | mean | std | min | 25% | 50% | 75% | max |
|---|---|---|---|---|---|---|---|---|
| siCD44 #1 | 3.0 | 0.08 | 0.01 | 0.07 | 0.075 | 0.08 | 0.085 | 0.09 |
| siCD44 #2 | 3.0 | 0.15 | 0.017 | 0.13 | 0.145 | 0.16 | 0.16 | 0.16 |
| siCtrl | 3.0 | 1.0 | 0.122 | 0.86 | 0.96 | 1.06 | 1.07 | 1.08 |

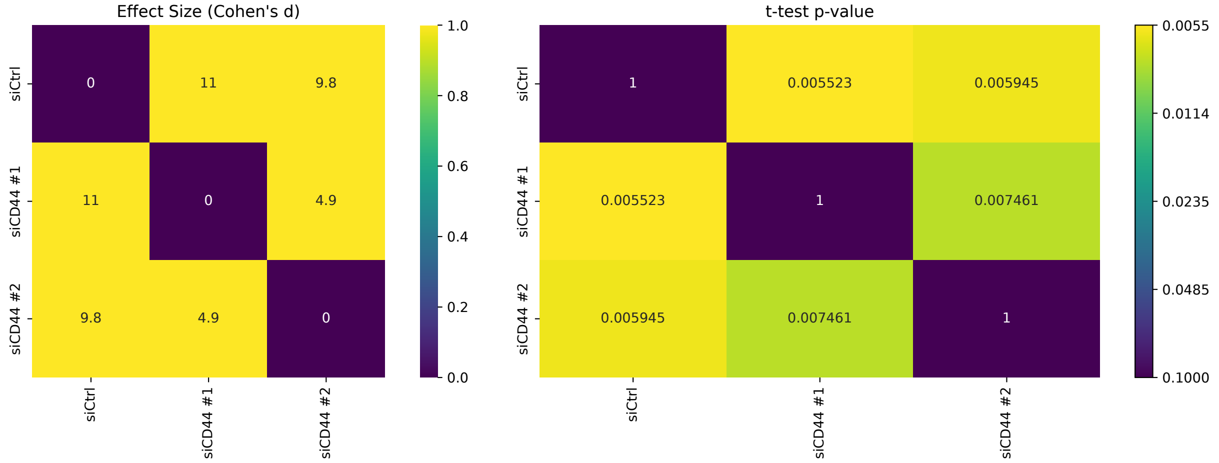

### FigS8A_MIAPaCa2

| Condition | count | mean | std | min | 25% | 50% | 75% | max |
|---|---|---|---|---|---|---|---|---|
| siCD44 #1 | 3.0 | 0.237 | 0.194 | 0.11 | 0.125 | 0.14 | 0.3 | 0.46 |
| siCD44 #2 | 3.0 | 0.41 | 0.114 | 0.33 | 0.345 | 0.36 | 0.45 | 0.54 |
| siCtrl | 3.0 | 1.0 | 0.03 | 0.97 | 0.985 | 1.0 | 1.015 | 1.03 |

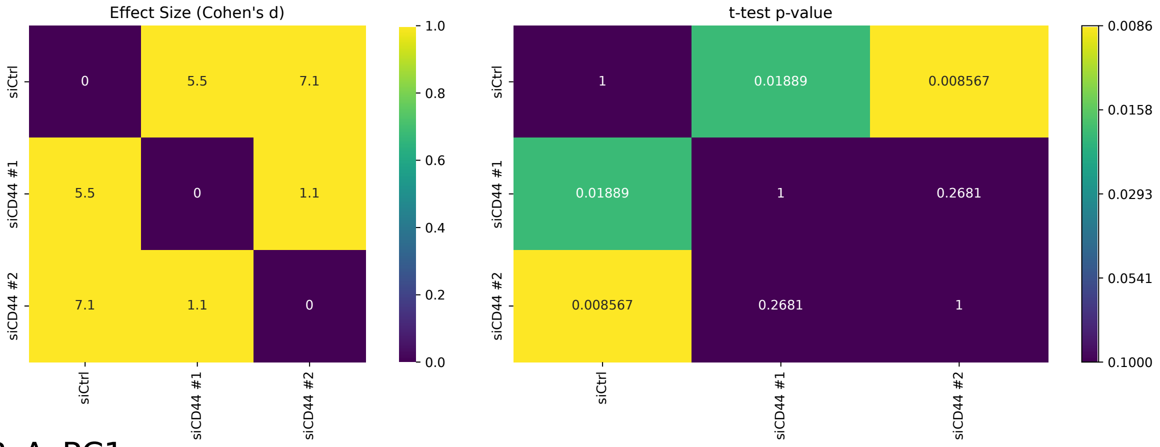


### FigS8B_AsPC1

| Condition | count | mean | std | min | 25% | 50% | 75% | max |
|---|---|---|---|---|---|---|---|---|
| siCD44 #1 | 3.0 | 0.237 | 0.194 | 0.11 | 0.125 | 0.14 | 0.3 | 0.46 |
| siCD44 #2 | 3.0 | 0.41 | 0.114 | 0.33 | 0.345 | 0.36 | 0.45 | 0.54 |
| siCtrl | 3.0 | 1.0 | 0.03 | 0.97 | 0.985 | 1.0 | 1.015 | 1.03 |

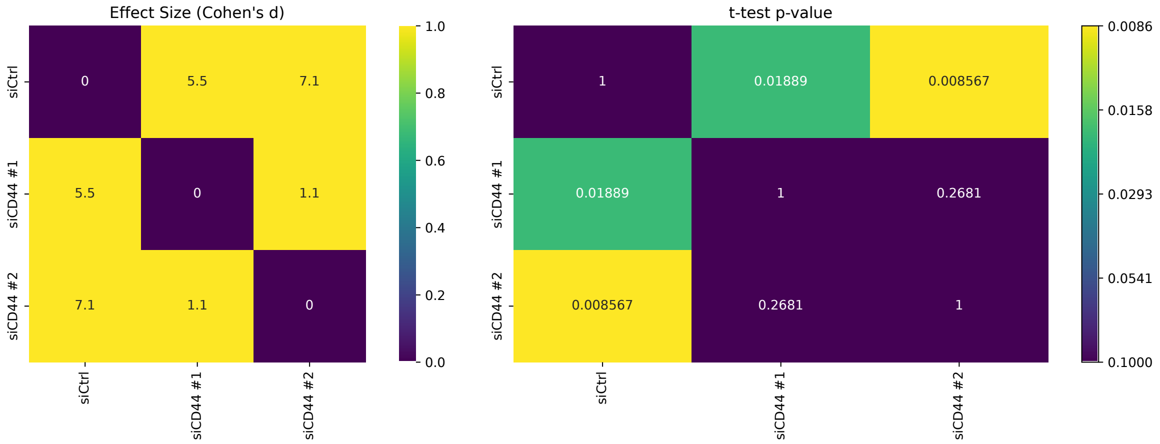

FigS8B_MIAPaCa2

| Condition | count | mean | std | min | 25% | 50% | 75% | max |
|---|---|---|---|---|---|---|---|---|
| siCD44 #1 | 4.0 | 0.062 | 0.079 | 0.013 | 0.018 | 0.027 | 0.07 | 0.179 |
| siCD44 #2 | 4.0 | 0.138 | 0.112 | 0.052 | 0.081 | 0.1 | 0.157 | 0.302 |
| siCtrl | 4.0 | 1.0 | 0.0 | 1.0 | 1.0 | 1.0 | 1.0 | 1.0 |

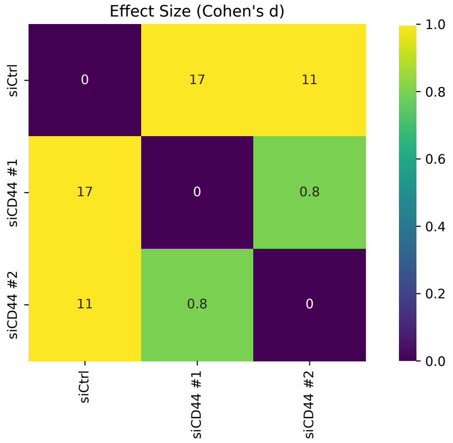


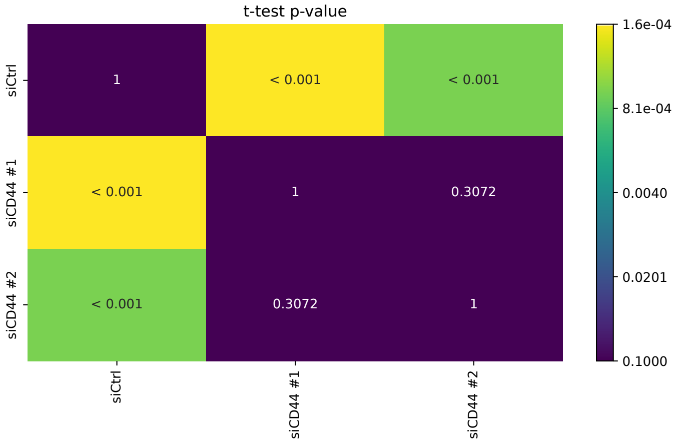


FigS8C_AsPC1

| Condition | count | mean | std | min | 25% | 50% | 75% | max |
|---|---|---|---|---|---|---|---|---|
| siCD44 #1 | 1385.0 | 0.712 | 0.399 | 0.0 | 0.453 | 0.634 | 0.865 | 4.363 |
| siCD44 #2 | 3124.0 | 0.475 | 0.262 | 0.0 | 0.322 | 0.429 | 0.57 | 3.995 |
| sictrl | 2451.0 | 1.0 | 0.573 | 0.0 | 0.6 | 0.901 | 1.284 | 5.665 |

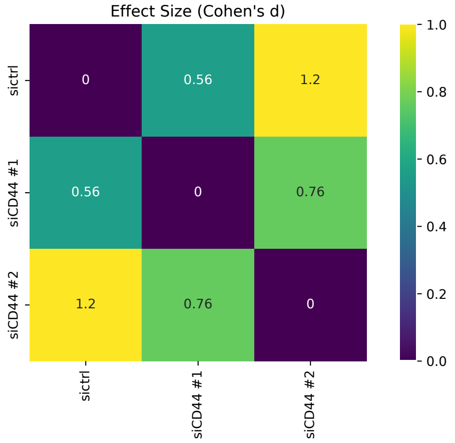


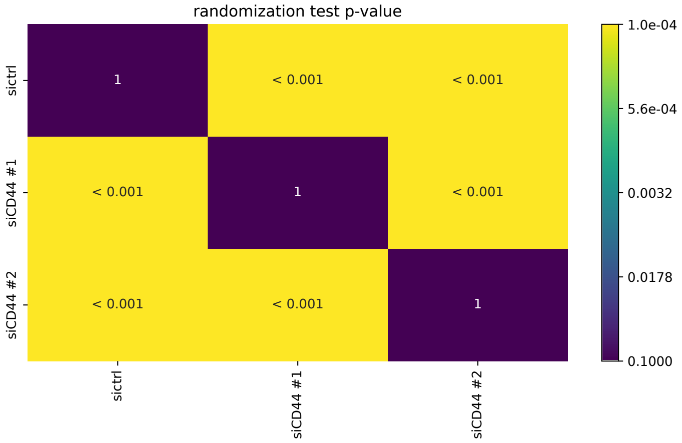

### FigS8C_MIAPaCa2

| Condition | count | mean | std | min | 25% | 50% | 75% | max |
|---|---|---|---|---|---|---|---|---|
| siCD44 #1 | 862.0 | 0.611 | 0.638 | 0.0 | 0.19 | 0.375 | 0.845 | 6.771 |
| siCD44 #2 | 199.0 | 0.43 | 0.212 | 0.0 | 0.321 | 0.411 | 0.566 | 1.182 |
| sictrl | 1230.0 | 1.0 | 0.517 | 0.0 | 0.642 | 0.914 | 1.27 | 4.459 |

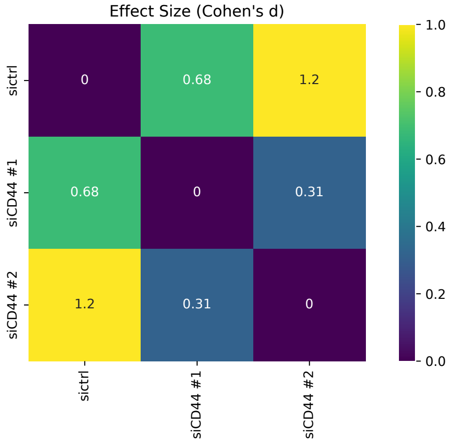


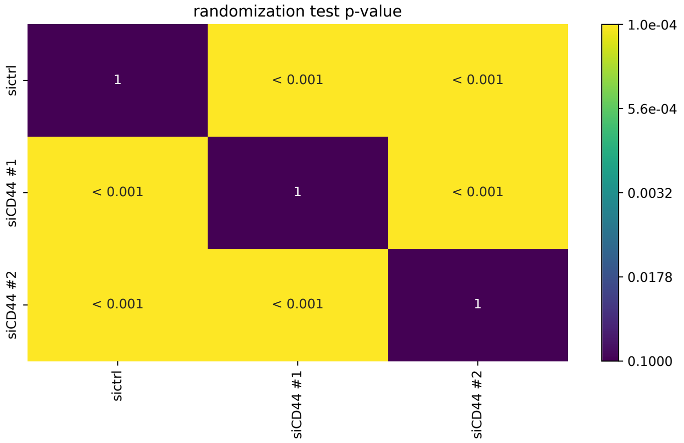

FigS8D

| Condition | count | mean | std | min | 25% | 50% | 75% | max |
|---|---|---|---|---|---|---|---|---|
| As 100 ctrl | 5242.0 | 485.773 | 408.206 | 5.486 | 147.746 | 375.665 | 713.576 | 1696.048 |
| As 100 si1 | 7772.0 | 926.57 | 415.711 | 10.689 | 548.913 | 995.124 | 1342.093 | 1640.448 |
| As 100 si2 | 5645.0 | 923.65 | 433.265 | 10.465 | 522.177 | 1023.331 | 1342.354 | 1727.117 |
| As 200 ctrl | 4723.0 | 611.024 | 460.587 | 7.199 | 169.76 | 541.12 | 1002.926 | 1791.801 |
| As 200 s1 | 9462.0 | 1081.967 | 311.123 | 15.89 | 829.232 | 1280.965 | 1335.378 | 1798.83 |
| As 200 s2 | 6158.0 | 1084.548 | 333.416 | 9.555 | 815.586 | 1327.552 | 1335.922 | 1664.166 |
| As 300si1 | 5871.0 | 1018.62 | 281.039 | 10.619 | 791.21 | 1024.617 | 1323.923 | 1545.27 |
| As 300 ctrl | 2166.0 | 678.813 | 437.269 | 9.966 | 228.252 | 712.28 | 1036.826 | 1439.882 |
| As 300 si2 | 2348.0 | 1027.08 | 336.936 | 10.795 | 803.964 | 1109.029 | 1328.384 | 1512.904 |
| As Wash ctrl | 5553.0 | 312.823 | 318.539 | 4.48 | 88.143 | 188.238 | 421.386 | 1677.915 |
| As wash si1 | 7400.0 | 956.386 | 301.477 | 16.313 | 722.691 | 943.473 | 1273.29 | 1709.416 |
| As wash si2 | 4315.0 | 1015.858 | 308.939 | 11.536 | 784.645 | 1042.032 | 1326.735 | 1577.229 |
| Mia 100 ctrl | 9137.0 | 259.401 | 228.17 | 2.144 | 86.002 | 194.878 | 382.097 | 2282.474 |
| Mia 100 si1 | 7832.0 | 1021.088 | 399.023 | 11.725 | 686.488 | 1307.045 | 1343.84 | 1549.84 |
| Mia 100 si2 | 9134.0 | 803.831 | 427.366 | 9.907 | 434.231 | 754.47 | 1333.81 | 1678.304 |
| Mia 200 ctrl | 7605.0 | 347.798 | 323.628 | 6.478 | 113.757 | 231.04 | 484.706 | 1822.957 |
| Mia 200 si1 | 11323.0 | 1111.654 | 313.066 | 12.07 | 887.935 | 1330.984 | 1335.96 | 1631.229 |
| Mia 200 si2 | 10263.0 | 971.06 | 344.323 | 12.747 | 672.526 | 989.61 | 1333.225 | 1722.317 |
| Mia 300 ctrl | 2306.0 | 501.848 | 415.114 | 9.641 | 140.759 | 348.577 | 815.924 | 1632.511 |
| Mia 300 si1 | 6284.0 | 1001.073 | 286.608 | 15.22 | 784.093 | 995.214 | 1320.344 | 1646.687 |
| Mia 300 si2 | 4040.0 | 927.81 | 316.348 | 15.309 | 743.447 | 916.471 | 1208.683 | 1599.858 |
| Mia wash ctrl | 15056.0 | 195.78 | 168.568 | 4.473 | 77.655 | 148.779 | 259.017 | 1446.188 |
| Mia wash si1 | 7895.0 | 1065.938 | 281.015 | 14.388 | 835.135 | 1136.119 | 1329.09 | 1849.005 |
| Mia wash si2 | 4629.0 | 915.572 | 337.258 | 11.703 | 704.421 | 899.142 | 1270.554 | 1524.129 |

FigS8E

| Condition | count | mean | std | min | 25% | 50% | 75% | max |
|---|---|---|---|---|---|---|---|---|
| As 100 ctrl | 5242.0 | 0.594 | 0.411 | -0.518 | 0.122 | 0.783 | 0.987 | 1.0 |
| As 100 si1 | 7772.0 | 0.965 | 0.134 | -0.368 | 0.984 | 0.993 | 0.996 | 1.0 |
| As 100 si2 | 5645.0 | 0.959 | 0.15 | -0.046 | 0.984 | 0.993 | 0.996 | 1.0 |
| As 200 ctrl | 4723.0 | 0.735 | 0.343 | -0.325 | 0.496 | 0.953 | 0.996 | 1.0 |
| As 200 s1 | 9462.0 | 0.986 | 0.075 | -0.201 | 0.993 | 0.998 | 0.999 | 1.0 |
| As 200 s2 | 6158.0 | 0.98 | 0.105 | -0.06 | 0.992 | 0.998 | 0.999 | 1.0 |
| As 300si1 | 5871.0 | 0.975 | 0.107 | -0.051 | 0.982 | 0.998 | 0.999 | 1.0 |
| As 300 ctrl | 2166.0 | 0.764 | 0.346 | -0.482 | 0.597 | 0.977 | 0.998 | 1.0 |
| As 300 si2 | 2348.0 | 0.94 | 0.212 | -0.26 | 0.985 | 0.998 | 0.999 | 1.0 |
| As Wash ctrl | 5553.0 | 0.395 | 0.369 | -0.452 | 0.058 | 0.281 | 0.735 | 1.0 |
| As wash si1 | 7400.0 | 0.96 | 0.143 | -0.111 | 0.977 | 0.996 | 0.999 | 1.0 |
| As wash si2 | 4315.0 | 0.961 | 0.157 | -0.235 | 0.984 | 0.998 | 0.999 | 1.0 |
| Mia 100 ctrl | 9137.0 | 0.329 | 0.344 | -0.591 | 0.022 | 0.196 | 0.618 | 1.0 |
| Mia 100 si1 | 7832.0 | 0.975 | 0.121 | -0.076 | 0.989 | 0.994 | 0.996 | 1.0 |
| Mia 100 si2 | 9134.0 | 0.936 | 0.171 | -0.089 | 0.961 | 0.987 | 0.994 | 1.0 |
| Mia 200 ctrl | 7605.0 | 0.515 | 0.357 | -0.544 | 0.171 | 0.533 | 0.869 | 1.0 |
| Mia 200 si1 | 11323.0 | 0.985 | 0.088 | -0.161 | 0.995 | 0.998 | 0.999 | 1.0 |
| Mia 200 si2 | 10263.0 | 0.971 | 0.104 | -0.401 | 0.977 | 0.996 | 0.998 | 1.0 |
| Mia 300 ctrl | 2306.0 | 0.654 | 0.352 | -0.283 | 0.337 | 0.777 | 0.983 | 1.0 |
| Mia 300 si1 | 6284.0 | 0.968 | 0.123 | -0.245 | 0.979 | 0.997 | 0.999 | 1.0 |
| Mia 300 si2 | 4040.0 | 0.932 | 0.198 | -0.385 | 0.969 | 0.989 | 0.999 | 1.0 |
| Mia wash ctrl | 15056.0 | 0.307 | 0.308 | -0.678 | 0.044 | 0.214 | 0.535 | 0.999 |
| Mia wash si1 | 7895.0 | 0.979 | 0.094 | -0.326 | 0.984 | 0.999 | 0.999 | 1.0 |
| Mia wash si2 | 4629.0 | 0.928 | 0.207 | -0.454 | 0.971 | 0.995 | 0.999 | 1.0 |

FigS8G

| Condition | count | mean | std | min | 25% | 50% | 75% | max |
|---|---|---|---|---|---|---|---|---|
| As 100 EC block | 4226.0 | 410.418 | 362.447 | 3.127 | 135.985 | 318.401 | 535.725 | 1695.48 |
| As 100 all cells | 4537.0 | 789.148 | 446.674 | 9.177 | 373.299 | 730.558 | 1339.336 | 1542.901 |
| As 100 ctrl | 4649.0 | 373.99 | 346.8 | 5.591 | 105.888 | 269.101 | 519.874 | 1579.833 |
| As 200 EC block | 4543.0 | 544.006 | 425.216 | 9.338 | 179.704 | 433.353 | 830.669 | 1705.777 |
| As 200 TC block | 10628.0 | 858.704 | 368.968 | 10.012 | 537.513 | 829.423 | 1270.601 | 1718.176 |
| As 200 all cells | 5237.0 | 933.485 | 362.703 | 8.986 | 612.435 | 942.623 | 1332.647 | 1476.521 |
| As 200 ctrl | 4948.0 | 495.152 | 434.997 | 8.912 | 120.614 | 349.262 | 789.508 | 1515.585 |
| As 300 EC block | 1953.0 | 713.747 | 415.581 | 7.827 | 383.995 | 734.469 | 1037.755 | 1389.029 |
| As 300 TC block | 4348.0 | 908.28 | 325.438 | 9.589 | 672.332 | 887.466 | 1235.502 | 1541.533 |
| As 300 all cells | 1173.0 | 842.91 | 349.861 | 6.98 | 612.17 | 835.074 | 1129.09 | 1362.19 |
| As 300 ctrl | 2133.0 | 616.643 | 414.277 | 8.841 | 196.063 | 631.405 | 915.714 | 1745.153 |
| As wash EC block | 7050.0 | 279.701 | 278.225 | 3.68 | 83.813 | 182.209 | 384.215 | 1719.056 |
| As wash TC block | 6796.0 | 852.436 | 319.36 | 12.054 | 611.292 | 817.714 | 1111.699 | 1863.259 |
| As wash all cells | 3807.0 | 916.407 | 344.203 | 9.263 | 623.092 | 912.668 | 1319.514 | 1541.387 |
| As wash ctrl | 5365.0 | 286.478 | 288.508 | 3.351 | 88.78 | 182.067 | 376.022 | 1556.392 |
| Mia 100 EC block | 6251.0 | 306.547 | 287.605 | 2.058 | 85.139 | 243.521 | 406.238 | 1650.559 |
| Mia 100 TC block | 11940.0 | 711.202 | 432.558 | 7.906 | 325.216 | 609.678 | 1140.829 | 1580.885 |
| Mia 100 all cells | 13558.0 | 571.887 | 378.546 | 8.674 | 281.271 | 460.39 | 795.869 | 1719.107 |
| Mia 100 ctrl | 7444.0 | 294.724 | 294.374 | 0.303 | 78.169 | 210.212 | 397.546 | 1615.223 |
| Mia 200 EC block | 5382.0 | 342.547 | 379.956 | 1.313 | 75.723 | 188.769 | 438.314 | 1600.355 |
| Mia 200 TC block | 15014.0 | 869.578 | 386.714 | 15.174 | 521.867 | 836.916 | 1332.334 | 1620.37 |
| Mia 200 all cells | 11392.0 | 792.91 | 400.733 | 11.329 | 442.535 | 727.933 | 1251.534 | 1749.462 |
| Mia 200 ctrl | 7684.0 | 324.979 | 343.808 | 1.564 | 82.797 | 189.055 | 439.604 | 1709.658 |
| Mia 300 EC block | 2758.0 | 605.454 | 484.749 | 11.075 | 124.05 | 565.677 | 1050.84 | 1635.223 |
| Mia 300 TC block | 5489.0 | 955.272 | 338.388 | 11.486 | 665.87 | 952.542 | 1328.827 | 1458.885 |
| Mia 300 all cells | 7529.0 | 867.847 | 318.119 | 13.921 | 618.228 | 816.633 | 1154.383 | 1590.61 |
| Mia 300 ctrl | 3209.0 | 479.307 | 417.805 | 9.27 | 120.305 | 308.423 | 762.032 | 1488.739 |
| Mia wash EC block | 7434.0 | 236.188 | 206.947 | 0.044 | 70.521 | 188.007 | 357.946 | 1657.799 |
| Mia wash TC block | 8281.0 | 975.304 | 354.144 | 9.869 | 679.379 | 1021.204 | 1331.639 | 1696.861 |
| Mia wash all cells | 9665.0 | 848.011 | 367.893 | 13.888 | 563.576 | 806.77 | 1242.407 | 1576.187 |
| Mia wash ctrl | 9814.0 | 223.599 | 242.993 | 2.121 | 67.397 | 142.089 | 290.971 | 1623.835 |
| As 100 TC block | 9078.0 | 697.593 | 411.416 | 10.733 | 348.076 | 583.04 | 1057.286 | 1672.948 |

FigS8G

| Condition | count | mean | std | min | 25% | 50% | 75% | max |
|---|---|---|---|---|---|---|---|---|
| As 100 EC block | 4226.0 | 0.467 | 0.425 | -0.674 | 0.024 | 0.394 | 0.956 | 1.0 |
| As 100 all cells | 4537.0 | 0.944 | 0.177 | -0.064 | 0.973 | 0.987 | 0.993 | 1.0 |
| As 100 ctrl | 4649.0 | 0.546 | 0.408 | -0.522 | 0.095 | 0.642 | 0.965 | 1.0 |
| As 200 EC block | 4543.0 | 0.692 | 0.367 | -0.266 | 0.366 | 0.915 | 0.992 | 1.0 |
| As 200 TC block | 10628.0 | 0.971 | 0.116 | -0.38 | 0.983 | 0.995 | 0.998 | 1.0 |
| As 200 all cells | 5237.0 | 0.981 | 0.09 | -0.257 | 0.988 | 0.996 | 0.998 | 1.0 |
| As 200 ctrl | 4948.0 | 0.693 | 0.345 | -0.395 | 0.424 | 0.86 | 0.987 | 1.0 |
| As 300 EC block | 1953.0 | 0.82 | 0.328 | -0.263 | 0.874 | 0.989 | 0.998 | 1.0 |
| As 300 TC block | 4348.0 | 0.957 | 0.166 | -0.391 | 0.983 | 0.997 | 0.999 | 1.0 |
| As 300 all cells | 1173.0 | 0.933 | 0.229 | -0.281 | 0.991 | 0.998 | 0.999 | 1.0 |
| As 300 ctrl | 2133.0 | 0.785 | 0.312 | -0.327 | 0.624 | 0.972 | 0.997 | 1.0 |
| As wash EC block | 7050.0 | 0.35 | 0.365 | -0.493 | 0.031 | 0.201 | 0.678 | 0.999 |
| As wash TC block | 6796.0 | 0.961 | 0.149 | -0.356 | 0.984 | 0.997 | 0.998 | 1.0 |
| As wash all cells | 3807.0 | 0.973 | 0.124 | -0.215 | 0.989 | 0.996 | 0.998 | 1.0 |
| As wash ctrl | 5365.0 | 0.437 | 0.372 | -0.654 | 0.088 | 0.363 | 0.815 | 1.0 |
| Mia 100 EC block | 6251.0 | 0.323 | 0.386 | -0.598 | 0.006 | 0.104 | 0.697 | 1.0 |
| Mia 100 TC block | 11940.0 | 0.936 | 0.162 | -0.542 | 0.957 | 0.984 | 0.993 | 1.0 |
| Mia 100 all cells | 13558.0 | 0.895 | 0.213 | -0.39 | 0.928 | 0.972 | 0.989 | 1.0 |
| Mia 100 ctrl | 7444.0 | 0.327 | 0.391 | -0.74 | 0.009 | 0.104 | 0.727 | 1.0 |
| Mia 200 EC block | 5382.0 | 0.455 | 0.388 | -0.476 | 0.079 | 0.374 | 0.886 | 1.0 |
| Mia 200 TC block | 15014.0 | 0.97 | 0.095 | -0.079 | 0.976 | 0.994 | 0.997 | 1.0 |
| Mia 200 all cells | 11392.0 | 0.947 | 0.146 | -0.184 | 0.964 | 0.993 | 0.997 | 1.0 |
| Mia 200 ctrl | 7684.0 | 0.501 | 0.369 | -0.529 | 0.134 | 0.503 | 0.877 | 1.0 |
| Mia 300 EC block | 2758.0 | 0.708 | 0.37 | -0.462 | 0.433 | 0.963 | 0.998 | 1.0 |
| Mia 300 TC block | 5489.0 | 0.974 | 0.115 | -0.033 | 0.987 | 0.998 | 0.999 | 1.0 |
| Mia 300 all cells | 7529.0 | 0.97 | 0.12 | -0.291 | 0.983 | 0.998 | 0.999 | 1.0 |
| Mia 300 ctrl | 3209.0 | 0.689 | 0.342 | -0.501 | 0.418 | 0.829 | 0.991 | 1.0 |
| Mia wash EC block | 7434.0 | 0.196 | 0.298 | -0.695 | 0.007 | 0.041 | 0.271 | 0.999 |
| Mia wash TC block | 8281.0 | 0.965 | 0.142 | -0.267 | 0.987 | 0.998 | 0.999 | 1.0 |
| Mia wash all cells | 9665.0 | 0.927 | 0.204 | -0.374 | 0.971 | 0.994 | 0.998 | 1.0 |
| Mia wash ctrl | 9814.0 | 0.31 | 0.341 | -0.573 | 0.028 | 0.157 | 0.584 | 1.0 |
| As 100 TC block | 9078.0 | 0.948 | 0.145 | -0.201 | 0.96 | 0.983 | 0.992 | 1.0 |
